# Inducible, but not constitutive, pancreatic *REG/Reg* isoforms are regulated by intestinal microbiota and pancreatic diseases

**DOI:** 10.1101/2024.10.18.619139

**Authors:** Yixuan Zhou, Macy R. Komnick, Fabiola Sepulveda, Grace Liu, Elida Nieves-Ortiz, Kelsey Meador, Ornella Ndatabaye, Aliia Fatkhullina, Natalie J. Wu-Woods, Paulina M. Naydenkov, Johnathan Kent, Nathaniel Christiansen, Maria L Madariaga, Piotr Witkowski, Rustem F. Ismagilov, Daria Esterházy

## Abstract

The *REG*/*Reg* gene locus encodes for a conserved family of potent antimicrobial but also pancreatitis-associated proteins. Here we investigated whether *REG/Reg* family members differ in their baseline expression levels and abilities to be regulated in the pancreas and gut upon perturbations. We found, in human and mouse, pancreas and gut differed in *REG*/*Reg* isoform levels and preferences, with duodenum most resembling the pancreas. Pancreatic acinar cells and intestinal enterocytes were the dominant REG producers. Intestinal symbiotic microbes regulated the expression of the same, select *Reg* members in gut and pancreas. These *Reg* members had the most STAT3-binding sites close to the transcription start sites and were partially IL-22 dependent. We thus categorized them as “inducible” and others as “constitutive”. Indeed, also in models of pancreatic-ductal adenocarcinoma and pancreatitis, only inducible *Reg* members were upregulated in pancreas. While intestinal *Reg* expression remained unchanged upon pancreatic perturbation, pancreatitis altered the microbial composition of the duodenum and feces shortly after disease onset. Our study reveals differential usage and regulation of *REG*/*Reg* isoforms as a mechanism for tissue-specific innate immunity, highlights the intimate connection of pancreas and duodenum, and implies a gut-to-pancreas communication axis resulting in a coordinated *Reg* response.

## Introduction

Barrier sites of the body have the challenging task of maintaining a symbiotic relationship with commensal organisms while effectively killing pathogens. A broad set of innate and adaptive immune cells has evolved to maintain this delicate equilibrium. At barrier sites, the cells directly interfacing with microbes, such as keratinocytes or gut epithelial cells, also possess antimicrobial activity. Though often requiring communication with hematopoietic immune cells, they substantially contribute to keeping the symbiotic microbiota in check and enhancing pathogen clearance.

In the gut epithelium, one mechanism is the secretion of antimicrobial peptides or small proteins including the C-type lectin family of REG proteins. In humans, this family includes *REG1A*, *REG1B* (*REG1* subfamily), *REG3A*, *REG3G* (REG3 subfamily) clustered on Chromosome 2, and *REG4* on Chromosome 1. In mice, it consists of *Reg1* and *Reg2* (*Reg1* subfamily), *Reg3a*, *Regb3*, *Reg3g*, and *Reg3d* (*Reg3* subfamily) clustered on Chromosome 6 and *Reg4* on Chromosome 3. Better studied in murine intestine, REG3γ was found to spatially restrain the microbiota from contacting the epithelium and target gram-positive bacteria^1,2^, while REG3β may bind both gram-positive and gram-negative^3^, and REG3A gram negative bacteria^4^. Mechanistically, human REG3A has been found to form filaments of hexameric, pore-forming rings *in vitro* that insert into bacterial membranes to kill^5^ and REG3G to aggregate bacteria^6^. *Reg3g* expression is regulated by microbial colonization and some bacterial infections^7–9^, particularly induced by flagellin in an IL-22 dependent manner^10^, and upregulated by elevated IL-6, another infection-associated cytokine^11^. Both cytokine signaling pathways converge on STAT3 activation. In humans, intestinal *REG* expression is elevated in many patients with inflammatory bowel diseases^12^ and colorectal cancer^13^, suggesting a link with the microbiota and inflammation.

Surprisingly, such a bactericidal role for REG proteins has scarcely been proposed in the pancreas^6^, even though the gene family was originally discovered there. REG proteins were initially observed to be upregulated upon pancreatitis (hence their alternative name, pancreatitis-associated protein, or PAP) and after partial pancreatectomy, where they were thought to aid beta cell regeneration^14^ (from which the name REG is derived). REG1A, REG1B and REG3A are present in human pancreatic juice even under non-disease conditions^15,16^. However, elevated pancreatic *REG* expression is associated with poor prognosis in pancreatic ductal adenocarcinoma (PDAC)^17,18^, and in mice overexpression of *Reg3g* is sufficient to aggravate PDAC in a genetic model^19^. Knocking out the entire pancreatic *Reg* family is protective in cerulein-induced pancreatitis^20^. Overall, a pro-proliferative role has been assigned to REG proteins in the pancreas^14^.

While C-type lectins can execute pleiotropic functions^21^, a potential bactericidal role of pancreatic REG proteins and their transcriptional regulation by microbes remain unclear. Additionally, most studies focusing on the gut, even if occasionally also analyzing the pancreas, measure only *Reg3g* and *Reg3b*, which may not be the genes most relevant in the pancreas. Therefore, in this study, we performed a holistic investigation on all *Reg* family members’ baseline expression profile in pancreas and gut and their abilities to be regulated upon intestinal and pancreatic perturbations.

## Results

### Mouse and human *Reg/REG* gene family members are structurally conserved but differ in their ligand binding domains and expression pattern between pancreas and gut

Most prior studies of REG proteins focused on the REG3 subfamily. To assess if other REG family members are likely to function similarly, we first took a bioinformatic approach. Amino acid sequence alignment of all murine and human REG isoforms revealed several conserved features (Fig. S1A): An inhibitory N-terminal prosegment followed by a trypsin cleavage site^22^ and characteristic features of C-type lectins^23–26^. The REG subgroups notably differed in the three loops forming the carbohydrate recognition domain (CRD): REG3, but not REG1 and REG4, subfamily members possess a longer loop 1 that includes the classical mannose binding motif EPN. REG1 and REG4 members are also more similar to each other than to REG3 proteins in the backloop, while REG1 proteins resemble REG3 sequences more in loop 2. Additional subtle differences within REG subfamilies exist in the CRD region. To understand how these primary sequences may translate into three-dimensional differences, we simulated the structure of all REG proteins using AlphaFold, noting that the software cannot account for structural shifts upon prosegment removal^22^ (Fig. S1B). The prosegment was not folded onto the proteins as predicted by NMR^22^, but the heads representing mature REG proteins were similar to the experimentally solved REG3A structure^27^ and predicted to display the canonical C-type lectin fold with two alpha helices “sandwiching” antiparallel beta sheets, whose loops cluster on one side of the protein to form the CRD ^23–26^. Notably, the CRD loops of the REG3 subfamily uniquely displayed one or two short beta strands and had a narrower binding cleft than subfamily REG1 and REG4. Scoring atomic distances within the REG proteins head regions (Fig. 1A) revealed structural similarity between members of the same REG subfamily regardless of species. The REG1 family was equally similar to the REG3 and REG4 subfamilies, while the REG3 and REG4 subfamilies were the most dissimilar. This was also reflected at the amino acid sequence similarity level (Fig. 1A, Fig. S1C). Overall, this analysis suggests that the REG subfamilies bind different ligands, and REG proteins within subfamilies may bind ligands with different affinities.

**Figure 1.**
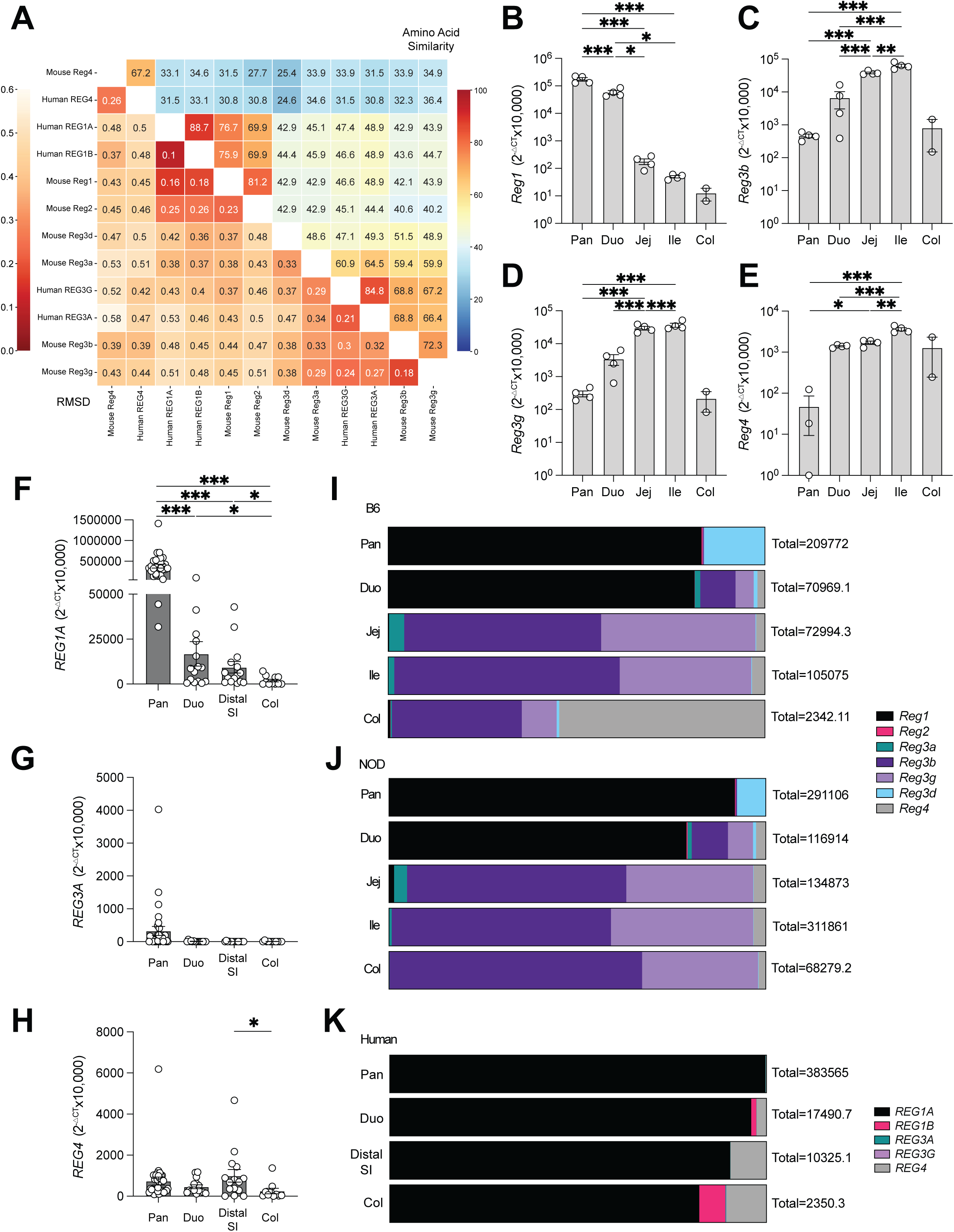
Mouse and human *Reg/REG* gene family members are structurally highly conserved but differ in their expression patterns between pancreas and gut. **A**. Sequence similarity (upper right corner) and structural similarity (lower left corner) of all mouse and human REG proteins excluding pro-segment regions. **B-E**. Expression levels of indicated *Reg* genes in the pancreas (pan), duodenum (duo), jejunum (jej), ileum (ile) and colon (col) of C57Bl6 mice by Q-PCR (*n*=2-4). **F-H**. Expression levels of indicated *REG* genes in human pancreas (pan, *n*=31), duodenum (duo, *n*=15), distal small intestine (distal SI, *n*=14), and colon (col, *n*=11) by Q-PCR. **I-K**. Relative contribution of *Reg/REG* isoforms to total *Reg/REG* pool in C57Bl6 mice (**I**), NOD mice (**J**) and humans (**K**). The numbers on right indicate total expression of *Reg/REG* genes. * p<0.05, ** p<0.01, *** p<0.001 by ANOVA.

Next, we compared isoform usage between the pancreas and different regions along the gut in B6 and NOD mice, as well as humans (Table 1). In both mouse strains, *Reg1*, *Reg2*, and *Reg3d* were more highly expressed in pancreas; *Reg3a* was highest in the jejunum, all other isoforms in the ileum. Within the gut, *Reg1*, *Reg2*, and *Reg3d* were most highly transcribed in the duodenum, while the colon consistently showed the lowest expression (Fig. 1B-E, Fig. S2A-J). The pancreas very lowly expressed *Reg4*, making it the most gut-specific isoform. Grossly, *Reg1* and *Reg3d* were the dominant pancreatic isoforms, *Reg1* and *Reg3b* were dominant in the duodenum, *Reg3b* and *Reg3g* in jejunum and ileum, and *Reg4* (B6 mice) or *Reg3b* and *Reg3*g (NOD mice) in the colon (Fig. 1I, J). The main difference between the mouse strains was that NOD mice had higher overall transcript levels, in line with previous reports on pancreas^28^. In humans the expression pattern was quite different. While *REG1* was highly expressed in the pancreas and displayed a proximal to distal gradient within the intestine (Fig. 1F), all other isoforms were lowly expressed despite robust pancreatic *CPA1* and intestinal *VILLIN1* expression, which rules out underrepresentation of acinar and gut epithelial cells, two cell types with previously reported *REG* expression^12,13,29^ (Fig. 1G, H; Fig. S2K-N). *REG4* was the second most highly expressed *REG* isoform in pancreas and gut, and no other isoform dominated the gut. Instead, the pancreas trended to exhibit the highest *REG3A* and *REG3G* expression. Overall, *REG1A* was the dominant isoform of both human pancreas and gut (Fig. 1K).

**Table 1.** Donor information of pancreas and gut samples.

These data suggest that the REG isoforms are utilized differently by the proximal gastrointestinal system (pancreas and duodenum) versus the distal gut, and are more tightly regulated in humans than mice, with the notable exception of *REG1A*.

### Gut enterocytes and pancreatic acinar cells are enriched for *REG*/*Reg* transcripts and the upstream signaling receptor *IL22RA1/Il22ra1*

We sought to more comprehensively understand how unique *REG/Reg* expression is to gut and pancreas at homeostasis, which cell types within these tissues express them, and how likely it was for *REG/Reg* to be activated by IL-22 and IL-6 ^10,30^ across tissues. We first measured the expression of all seven mouse *Reg* isoforms in a 22-tissue panel from B6 mice, including other important barrier sites like lung, skin and bladder (Fig. 2A). In this panel, only the pancreas and gut expressed *Reg*. To determine if the confinement of *Reg* expression to the pancreas and gut correlated with IL-22 and/or IL-6 receptor expression, we measured the transcript abundance of the subunits of IL-6 receptor (*Il6r1* and *Il6st*) and IL-22 receptor (*Il22ra1* and *Il10rb*) in our tissue panel, along with the cytokines themselves (*Il6*, *Il22*) and IL-22BP, a soluble IL-22 decoy receptor that sequesters IL-22 (*Il22ra2*)^31^ (Fig. 2B). *Il6r1* and *Il6st* were barely expressed in the gut; the pancreas displayed some expression, but similar levels were observed in organs that do not express *Reg* genes. By contrast, *Il22ra1* was enriched in the pancreas and small intestine (Fig. 2B, C). In addition, *Il22ra2* was enriched in the colon compared to pancreas and small intestine (Fig. 2B, C), correlating with the lower expression of *Reg* genes in the colon. *Il6* and *Il22* transcripts were very low, as typical for cytokines in whole tissue extracts, and their measurement was therefore omitted for the rest of this study. In human pancreas and gut, we found that *IL22R1* and *IL6ST* displayed higher transcript levels in the pancreas than gut and *IL22RA2* was the highest in the colon (Fig. 2D-G; Fig. S2O).

**Figure 2.**
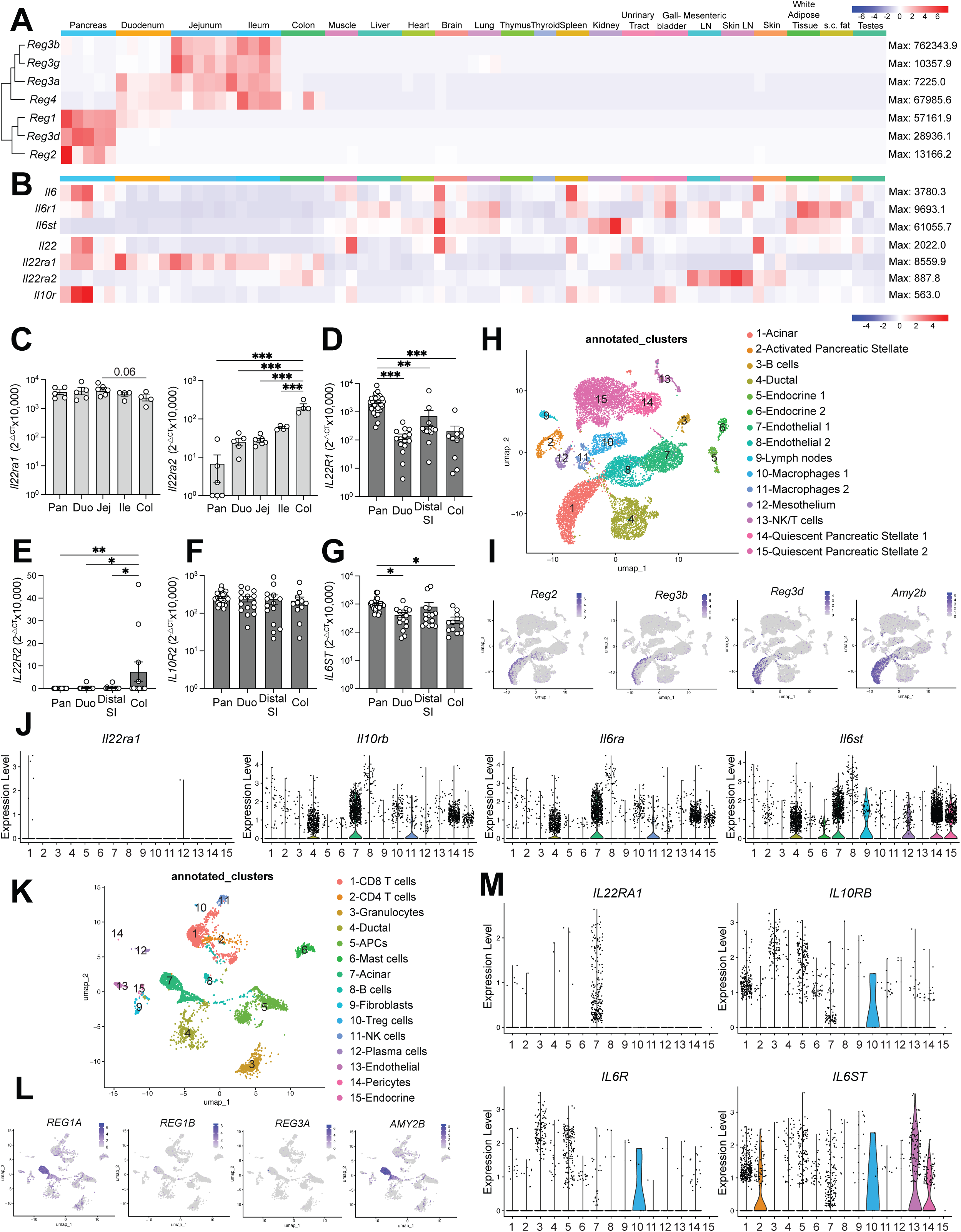
Gut and pancreas are enriched for *Reg* transcripts and the upstream signaling receptors. **A, B**. Heatmap of *Reg* gene (**A**) and IL-6 or IL-22 receptor signaling gene (**B**) expression in indicated tissues from 8-week-old C57Bl6 mice measured by Q-PCR (*n*=2-5, see as indicated by box graph). Maximal expression level per row is indicated at the right of each row. **C.** Expression of *Il22ra1* and *Il22ra2* in pancreas (pan), duodenum (duo), jejunum (jej), ileum (ile) and colon (col) of C57Bl6 mice by Q-PCR (*n*=4). **D-G**. Expression levels of indicated IL6 or IL22 receptor signaling genes in human pancreas (pan, *n*=31), duodenum (duo, *n*=15), distal small intestine (distal SI, n=14), and colon (col, *n*=11) by Q-PCR. **H**. Annotated *scRNAseq* feature plot of mouse pancreas. **I**. Feature plots of mouse *Reg2, Reg3b, Reg3d* and *Amy2b.* **J**. Violin plots of IL-6 or IL-22 receptor subunit genes across mouse pancreatic cell types. **K**. Annotated *scRNAseq* feature plot of human pancreas. **L**. Feature plots of human *REG1A*, *REG1B*, *REG3A* and *AMY2B*. **M**. Violin plot of IL-6 or IL-22 receptor subunit genes across human pancreatic cell types. * p<0.05, ** p<0.01, *** p<0.001 by ANOVA.

To pinpoint the cell types within pancreas and gut that express *REG*/*Reg* and assess whether this coincides with the expression of the IL22 receptor subunits, we re-analyzed published *scRNAseq* datasets. In mouse pancreas^32^, the expression of *Reg* isoforms was largely restricted to acinar cells (Fig. 2H, I), displaying a pattern on a feature plot similar to the acinar cell marker *Amy2b*. The *Il22ra1* signal recovery was low but confined to acinar cells (Fig. 2J). By contrast, *Il10rb*, *Il6r1, Il6st* (Fig. 2J), as well as *Il17ra* and *Il17rc,* the receptors for IL-17, which is often co-secreted with IL-22 (Fig. S3A), were co-enriched in ductal, endothelial, and pancreatic stellate cells. Similarly, in human pancreas^33^ *REG* and *IL22RA1* expression were confined to acinar cells (Fig. 2K-M).

*REG1A* was the dominant pancreatic *REG* isoform (Fig. 2L), confirming our Q-PCR results. Ductal and acinar cells expressed comparable levels of *Il10RB*, *Il6R, Il6ST* (Fig. 2M), while *Il17RA* and *Il17RC* were enriched in ductal compared to acinar cells (Fig. S3B). In neither the mouse nor human *scRNAseq* did we observe noteworthy expression of the receptor genes in pancreatic endocrine cells, though this could in part be due to the low representation of these cells in the datasets.

For murine intestine, we analyzed a *scRNAseq* dataset that encompassed epithelial cells from different regions of the small intestine^34^ (Fig. S3C, D). Feature plots for single genes (Fig. S3E) and their relation to cell types and region (Fig. S3C, D) showed that *Reg1* expression was enriched in duodenal enterocytes. *Reg2* and *Reg3d* were barely expressed. *Reg3a*, *Reg3b* and *Reg3g* were enriched in enterocytes of all small intestinal regions, though lower levels of *Reg3b* and *Reg3g* also mapped to virtually all cell types, including stem cells. *Reg4* was distinct, confined to goblet, Paneth, and enteroendocrine cells. Regarding cytokine receptor enrichment (Fig. S3F), *Il22ra1* and *Il17ra* were lowly expressed by nearly all cell types; *Il6ra* was enriched in endocrine cells. In human intestine, the scRNAseq dataset covered epithelial cells from the small and large intestine^35^ (Fig. S3G, H). *REG1A* was the most abundant isoform, predominantly expressed by small intestinal epithelial cells, with much lower levels in large intestine (Fig. S3I). Small intestinal enterocytes were the dominant population expressing *REG1A* and *REG1B*; *REG3A* and *REG3G* were also expressed by small intestinal stem cells, and *REG4* was enriched in goblet, Paneth and enteroendocrine cells. For cytokine receptors (Fig. S3J), *IL22RA1* was again enriched in small intestinal enterocytes, with noteworthy expression in small intestinal transit amplifying, stem and goblet cells, and large intestinal enterocytes. Other receptors displayed broader distribution among the 17 cell types, though enterocytes were the most frequent cell type.

Taken together, these data identify small intestinal enterocytes and pancreatic acinar cells as the major cell types expressing most *REG/Reg* isoforms under non-disease conditions.

### Pancreatic and intestinal *Reg* gene family transcripts correlate and are regulated by microbial colonization

The strong correlation of *REG/Reg* expression with that of *IL22RA1/Il22ra1*, but not IL-6 receptor subunits, suggested that IL-22 but not IL-6 contributes to baseline *REG/Reg* expression, though it could alternatively merely indicate a unique sensitivity of the pancreas and gut to IL-22. To determine whether the higher baseline expression of *REG/Reg* genes in pancreas and gut is driven by IL-22 as well as the gut microbiota, we first measured *Reg* gene expressions in pancreas, duodenum and colon of wildtype, heterozygous and *Il22* knockout mice (Fig. 3A, B, Fig. S4A). The duodenum showed the greatest sensitivity to IL-22 deficiency, with a 5-fold decrease in *Reg3a*, *Reg3b*, *Reg3g* and a downward trend in *Reg2* expression in knockout mice compared to wildtype. Similar but less pronounced trends were observed in pancreas and colon. By contrast, *Reg1*, *Reg3d* and *Reg4* were unaffected. These findings suggest that while IL-22 contributes to baseline *Reg* expression of 4 isoforms in SPF mice, other, IL-22-independent mechanisms, must contribute to baseline *Reg* expression, especially for *Reg1*, *Reg3d* and *Reg4*. To address the involvement of microbiota, we treated adult mice with broad-spectrum antibiotics. This resulted in lower expression of *Reg2, Reg3a and Reg3b* in pancreas, with *Reg1* and *Reg3g* trending down (Fig. 3C). Although antibiotics had no impact on duodenal *Reg* levels, it reduced *Reg3b, Reg3g* and *Reg3d* expression in the ileum (Fig. 3D, E). Conversely, in comparing the pancreata of germ free (GF) mice and ex-GF (F2 generation of GF mice colonized with Jax microbiota) mice, we observed an upregulation of *Reg1* and *Reg3g* in ex-GF mice (Fig. S4B-D). To understand how early in life baseline *Reg* expression is established, we measured *Reg* expression in pancreas of one-day-old (P1, pre-microbial colonization) and 28-day-old (P28, post-microbial colonization) GF and ex-GF mice. P1 mice already exhibited robust *Reg1* expression independently of the microbiota (Fig. S4E), similarly to another unrelated bacteriostatic protein, *Gp2* (Fig. S4J). Other *Reg* genes displayed low expression at P1, but increased by P28 in ex-GF but not GF mice, though *Reg2* also trended upward in GF mice (Fig. S4F-I).

**Figure 3.**
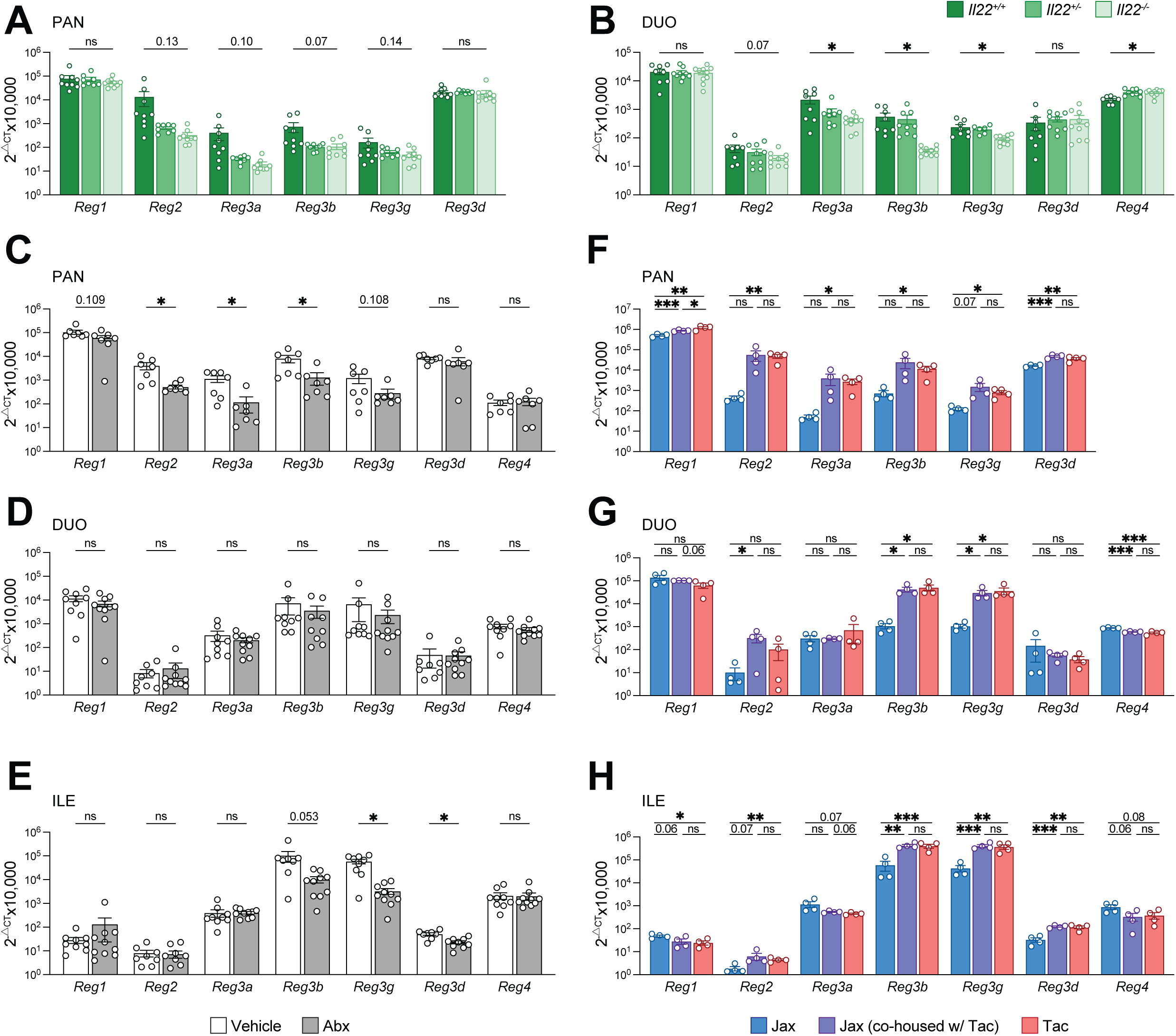
Pancreatic and intestinal *Reg* gene family transcripts correlate and are regulated by microbial colonization. **A-C.** Expression levels of indicated *Reg* genes in pancreas, duodenum and colon of *Il22*^+/+^, *Il22*^+/-^ and *Il22*^−/−^ mice (n=9, 9 and 10, respectively) by Q-PCR. Data pooled from two independent experiments. **D-I**. Pancreatic expressions of indicated *Reg* genes in germ-free (GF) or housing matched specific pathogen free (exGF) C57Bl6 mice (*n*=3), or C57Bl6 mice purchased from JAX, or Taconic (Tac), or JAX-mice co-housed with Taconic mice (*n*=4) for 4 weeks prior to analysis by Q-PCR. **J-P**. Expression levels of indicated *Reg* genes in duodenum and ileum of GF and exGF mice (*n*=4), C57Bl6 mice purchased from JAX, or Taconic (Tac), or JAX-mice co-housed with Taconic mice (*n*=4) for 4 weeks prior to analysis by Q-PCR. Data on graphs E-J represent samples run on the same Q-PCR plate to permit direct comparison of expression levels between the two experiments. Representative data of two independent experiments is shown. * p<0.05, ** p<0.01, *** p<0.001 by student t-test.

The complex Jax microbiota likely increased pancreatic and intestinal *Reg* expression compared to GF mice through multiple IL-22 augmenting pathways, including increased abundance of microbe-dependent metabolites^9,36^ and PAMPs like flagellin^10,37^ that induce IL-23, IL-6, IL-1b and TGFb^10,11,30,38^ or direct ligation of transcription factors (e.g. AhR, RORgt), which contribute to the differentiation of IL-22 producers (ILC3, Th17, TCRgd, NK cells and neutrophils)^7,39^ or act on pre-existing tissue-resident immune cells. Additional IL-22 independent mechanisms by which microbes increase *REG/Reg* expression also exist^30,40,41^, though they often converge on STAT3 activation. Given that microbiota composition impacts how strongly these pathways are activated, we explored whether a microbiota enriched in SFB^42^, an ileum-resident, epithelium attaching, non-disseminating commensal known to rapidly induce IL-22 secretion by ILC3^38^ and later by Th17 cells^42^, leads to a pancreatic *Reg* response in concert with the already reported intestinal upregulation of *Reg3g* and *Reg3b*. We compared the expression of all *Reg* family members in the pancreas, duodenum and ileum of age-matched B6 mice from Jax (SFB-free) versus mice from Taconic (SFB-bearing)^42^ and Jax mice co-housed Taconic mice for 4 weeks to account for the slight genetic difference between the B6 strains. All *Reg* genes were more expressed in the pancreas of Taconic than Jax mice (Fig. 3F), typically by over 10-fold, except for *Reg1* and *Reg3d*, which were about 3-fold higher. Jax mice co-housed with Taconic mice elevated their pancreatic *Reg* levels to resemble those of Taconic mice (Fig. 3F). Exposure to the Taconic microbiota in the gut led to a more select upregulation of *Reg* isoforms (Fig. 3G, H). *Reg2*, *Reg3b*, *Reg3g* increased in both duodenum and ileum while *Reg3d* increased only in ileum. When looking for the common denominator of all these experiments (Fig.3, Fig. S4), it became evident that *Reg2, Reg3a, Reg3b* and *Reg3g* expression was more influenced by the microbiota and dependent on IL-22 in both gut and pancreas than *Reg1, Reg3d* and *Reg4*. This was in line with transcript factor binding site analysis showing that the more affected *Reg* genes have more STAT3 binding sites near the transcription start sites (Fig. S5A). Therefore, we categorized these highly regulated *Reg* genes as “inducible” and the others as “constitutive”. Notably, in humans, REG1A, which is highly expressed at baseline, has 3 STAT3-binding sites. We thus predict it to be more regulated by environmental factors than its mouse counterpart (Fig. S5B).

All in all, these data strongly support the idea that intestinal microbial colonization can selectively co-regulate the expression of inducible *Reg* genes in the pancreas and gut.

### Murine PDAC triggers *Reg* upregulation in pancreas but not gut

Poor prognosis in PDAC has been linked to increased *REG*3A and IL-22 levels^17,43^, though it is not clear whether this reflects the presence of a disease aggravating microbiota or is an independent feature of aggressive disease. To determine whether “primary pancreatic” diseases regulate *Reg* genes in pancreas and gut, we first utilized an inducible murine PDAC model with *Ptf1a^CreERT2^* mice on a *p53^fl/fl^* and *Kras^G12D^* background (“KPC” mice). In the pancreas, all *Reg* isoforms were upregulated in both the tumor and non-tumor tissues compared to controls as early as 4 weeks post-induction (Fig. 4A, B; Fig. S6A-E), when lesions start to appear (Fig. S6A). Notably, inducible *Reg* forms were the most drastically affected, increasing by almost 100-fold, while constitutive *Reg* forms, *Reg1* and *Reg3d*, were less impacted. Since *Reg3b* and *Reg3g* were strongly upregulated in the pancreas, we measured these genes in the gut as proxies for querying pancreas-to-gut regulation. However, even at 12 weeks, when the impact on these genes was the strongest in the pancreas (Fig. 4A, B, “non-tumor”), they were unchanged in the gut (Fig. 4C, D). This suggested that the mechanism upregulating *Reg* in primary PDAC stays local and is not transferred to the gut.

**Figure 4.**
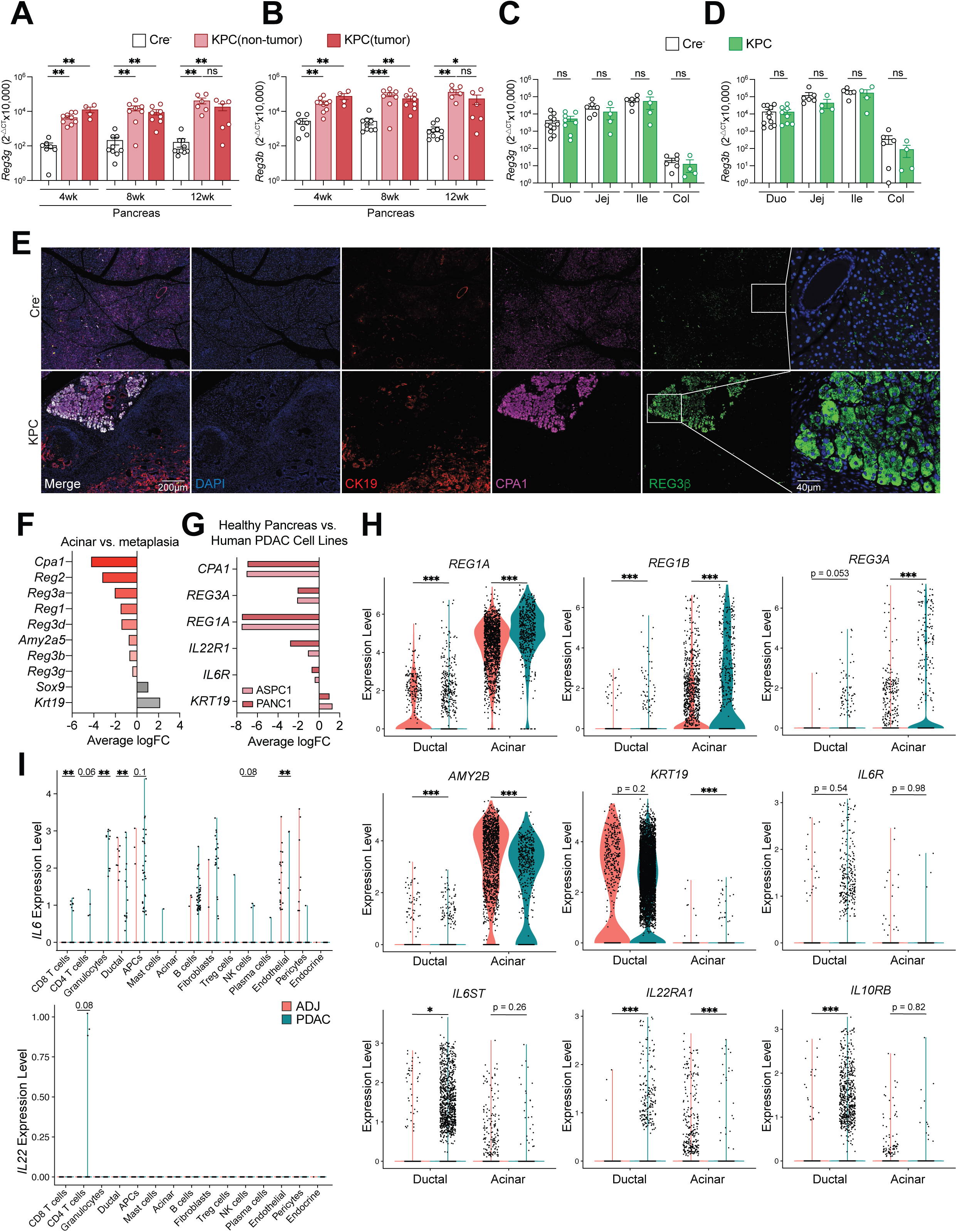
A mouse model of primary pancreatic ductal adenocarcinoma triggers *Reg* upregulation in pancreas but not gut. **A, B.** Pancreatic expressions of *Reg3g* (**A**) and *Reg3b* (**B**) in KPC mice or their Cre^−^ littermates at 4, 8 or 12 weeks post tamoxifen-induced PDAC onset measured by Q-PCR (WT *n*=7, 9,10, PDAC *n*=9, 8, 6 for 4, 8, 12 weeks, respectively). Data pooled from 3 cohorts per time point. **C, D.** Intestinal expressions of *Reg3g* (**C**) and *Reg3b* (**D**) in different gut segments of KPC mice or their Cre^−^ littermates at 12 weeks post tamoxifen-induced PDAC onset measured by Q-PCR (WT *n*=6 (Jej, Ile, Col) or 11 (Duo), PDAC *n*=4 (Jej, Ile, Col) or 8(Duo)). Data pooled from 2-3 independent cohorts. **E**. Pancreatic sections from KPC mice at 12 weeks post tamoxifen treatment or Cre^−^ littermates stained for ductal marker CK19, acinar marker CPA1, REG3β and DAPI (nuclei). Bar=200 µm, inset bar=40 µm. **F.** Bar graph of Log2 fold change of indicated genes in acinar versus metaplastic pancreatic cells in mouse *scRNAseq*. **G.** Bar graph of Log2 fold change of indicated genes in healthy human pancreas versus human PDAC cell lines analyzed by Q-PCR. **H**. Violin plots of *REG1A, REG1B, REG3A, AMY2B, KRT19, IL6R, IL6ST, IL22RA1* and *IL10RB* expressions in human acinar versus ductal cells in PDAC or adjacent tissues based on *scRNAseq.* **I.** Violin plots of *Il6* and *Il22* in all cell types recovered from human PDAC or healthy adjacent tissues analyzed by *scRNAseq*. * p<0.05, ** p<0.01, *** p<0.001 by student t-test.

Additionally, we noticed that from week 8 onward all isoforms exhibited lower *Reg* levels in the “tumor” tissue than “non-tumor” tissue. Likely, as acinar cells increasingly lose their identity, they also lose *Reg* expression. These trends correlated with the loss of acinar markers *Cpa1* and *Ptf1a* and the gain of ductal markers *Krt19*, *Sox9* and *Car2* (Fig. S6F-J). Similarly, while most IL-6 or IL-22 receptor subunits remained unchanged in the pancreas of KPC mice (Fig. S6K-M), *Il22ra1* decreased in the tumor samples from week 8 (Fig. S6M), potentially reflecting the loss of acinar identity. To reconcile the finding that *Reg* in whole tissue extracts went up with a potential loss in their expression in advanced tumors, we took several approaches. First, we co-stained pancreatic sections of KPC mice and their littermate controls with antibodies against mouse REG3β, acinar marker CPA1 and ductal marker CK19 (Fig. 4E). The staining revealed both a much stronger staining intensity of REG3β on KPC pancreas and an exclusion of REG3β in the transformed, CK19-positive areas. We then queried a publicly available RNAseq dataset comparing acinar versus metaplastic cells from KPC mice^44^ (Fig. 4F). All *Reg* genes were downregulated in metaplastic cells within the range of *Cpa1* and *Amy2a5* de-enrichment. To explore if this finding may be true in human PDAC as well, we compared the expression of *REG* genes and *IL22R1* in two human PDAC cell lines to that of healthy human pancreas (Fig. 4G), and found those genes downregulated while *KRT19* was upregulated. Additionally, we analyzed a publicly available scRNAseq dataset composed of PDAC tissue from 17 patients and 3 adjacent control tissues (ADJ)^33^ (Fig. S7A, B). When focusing on ductal and acinar cells from PDAC and ADJ (Fig. 4H),

*REG1A*, *REG1B* and *REG3A* were more highly expressed in acinar cells from PDAC tissue. Though acinar cell remained the major source of *REG* transcripts, the genes were also more highly expressed in the ductal cells of PDAC tissue, possibly due to a fraction of ductal cells being of acinar origin. To pursue this possibility, we generated a non-integrated UMAP of only the ductal and acinar cells in the dataset (Fig. S7C). Ductal cells were predominantly contributed by PDAC samples (Fig. S7D, E). When mapping *REG* genes along with *AMY2B* and *KRT19* onto this UMAP (Fig. S7F), it became clear that there was a zone of intermediate *REG* expression coinciding with a decrease in *AMYB2B* and gain of *KRT19*. Pseudotime analysis (Fig. S7G) confirmed a trajectory from acinar clusters 1,4 and 10 (Fig. S7H, I) to ductal clusters 0, 2,3, 5, 7-12, 14, 15 and 17 passing through acinar-ductal hybrid clusters 6, 13, 19 and 20, matching decreasing *REG1A* expression (Fig. S7F). *IL22RA1* was lost in acinar and gained relatively in ductal cells in PDAC tissue, and IL-6 receptor subunit *IL6ST* were also upregulated in PDAC ducts (Fig. 4H), phenomena again pointing to an acinar origin of some “ductal” cells. This dataset also included some immune cell populations. Analysis of IL-6 and IL-22 producers (Fig. 4I) revealed an enrichment of antigen presenting cells (APCs), fibroblasts, B cells and granulocytes expressing *Il6*, as well as IL-22-producing CD4 T cells in PDAC tissues. This suggests that both IL-6 and IL-22 contribute to *REG* upregulation in human PDAC. Overall, the mouse and human data support a model in which *REG/Reg* genes are initially upregulated in acinar cells of PDAC, but they gradually lose the expression during acinar-to-ductal transformation (Fig. S6N).

Finally, we asked whether changes in pancreatic *Reg* expression in KPC mice affected the gut microbiome. We hypothesized that duodenal microbiome, either mucosa-associated or luminal, might be the most sensitive. However, 16S rRNA gene sequencing showed no difference between the duodenal microbiota of cancerous mice and their littermates at week 12 according to principal component analysis (PCA, Fig. S7J, K). Total bacterial load was also unchanged in duodenal mucosa or lumen. Fecal microbiome composition was also unaltered, as was the pancreatic microbiome, where both bacterial and fungal loads were very low (*data not shown*). Some variation was due to cage effects^45,46^ (Fig. S7J, K, bottom) but even within cages mice did not differ, though we could not exclude potential coprophagy^47^ counteracting any impact of PDAC on the gut microbiome.

In sum, primary pancreatic acinar-to-ductal transformation is sufficient to significantly upregulates inducible *Reg* genes, with a smaller effect on constitutive *Reg* genes, in PDAC without affecting intestinal *Reg* expression. In human, PDAC acinar cells also upregulate *REG* genes. While PDAC progression in mice heavily depends on the presence of a microbiota^48,49^ and IL-22^43^, our data show that an intestinal dysbiosis is not required as a triggering event for *Reg* upregulation in PDAC, nor does PDAC lead to a drastic gut microbial shift.

### Cerulein-induced pancreatitis triggers upregulation of *Reg* genes in pancreas and modifies the duodenal microbiome

Pancreatitis, another major exocrine pancreatic disease, is a known context in which REG3A is upregulated^6^. Chronic pancreatitis also poses a risk factor for PDAC development^50,51^. We therefore interrogated the *Reg* regulation pattern in the context of cerulein-induced pancreatitis. The pancreas displays enlarged acinar cells and signs of fibrosis as early as one week after cerulein treatment commencement (Fig. S8A). After 1 week of treatment, inducible *Reg* genes, *Reg2*, *Reg3a*, *Reg3b* and *Reg3g,* were 25- to 100-fold upregulated (Fig. 5A, S8B), while *Reg3d* was only doubled after 4 weeks and *Reg1* remained unchanged until downregulated at 8 weeks. IL-6 receptor subunits were upregulated from 4 weeks of treatment, suggesting that like in human PDAC, IL-6 contributes to *Reg* upregulation in pancreatitis. By contrast, *Il22ra1* was downregulated after 8 weeks. REG3b protein staining confirmed its restriction to acinar cells (Fig. S8E). Analysis of a publicly available scRNAseq dataset^32^ (Fig. S8F, G) confirmed the upregulation of *Reg3g* and *Reg2* in acinar cells following acute cerulein treatment (Fig. 5C), though *Reg3d* was downregulated in this dataset. To test if the downregulation of *Reg1* and *Reg3d* (the two highest expressor at baseline) and *Il22ra1* after 8 weeks could be due to de-differentiation, similar to PDAC (Fig. S6N), we examined *Amy2b* and *Krt19* regulation in the scRNAseq dataset and by Q-PCR along acinar and ductal markers (Fig. S8D). Acinar cells showed decreased *Amy2b* and increased *Krt19* expression (Fig. 5C), and in total pancreatic extracts, acinar marker *Cpa1* was downregulated while ductal markers *Sox9* and *Car2* were upregulated (Fig. S8D). Globally, the pancreas underwent substantial cellular redistribution following cerulein treatment (Fig. S8F, G). Despite the lack of access to human pancreatitis samples, we re-examined our pancreatic donor cohort (Table 1) and noticed potential splits by smoking status, cardiovascular disease (CVD), age over 50 and alcohol use, all of which have been proposed as risk factors for pancreatitis^52,53^. *REG1A* and *REG4* levels were elevated in the CVD, smoking and age over 50 donors (Fig. S2P-Q). Alcohol consumption had minimal impact (*data not shown*), possibly due to moderate use. However, increased expression of other acinar markers, like *CPA1*, *MIST1* (Fig. S2R-S) and *PTF1A* (*data not shown*), also co-segregated with these factors, indicating a more generic acinar renewal program may be at play.

**Figure 5.**
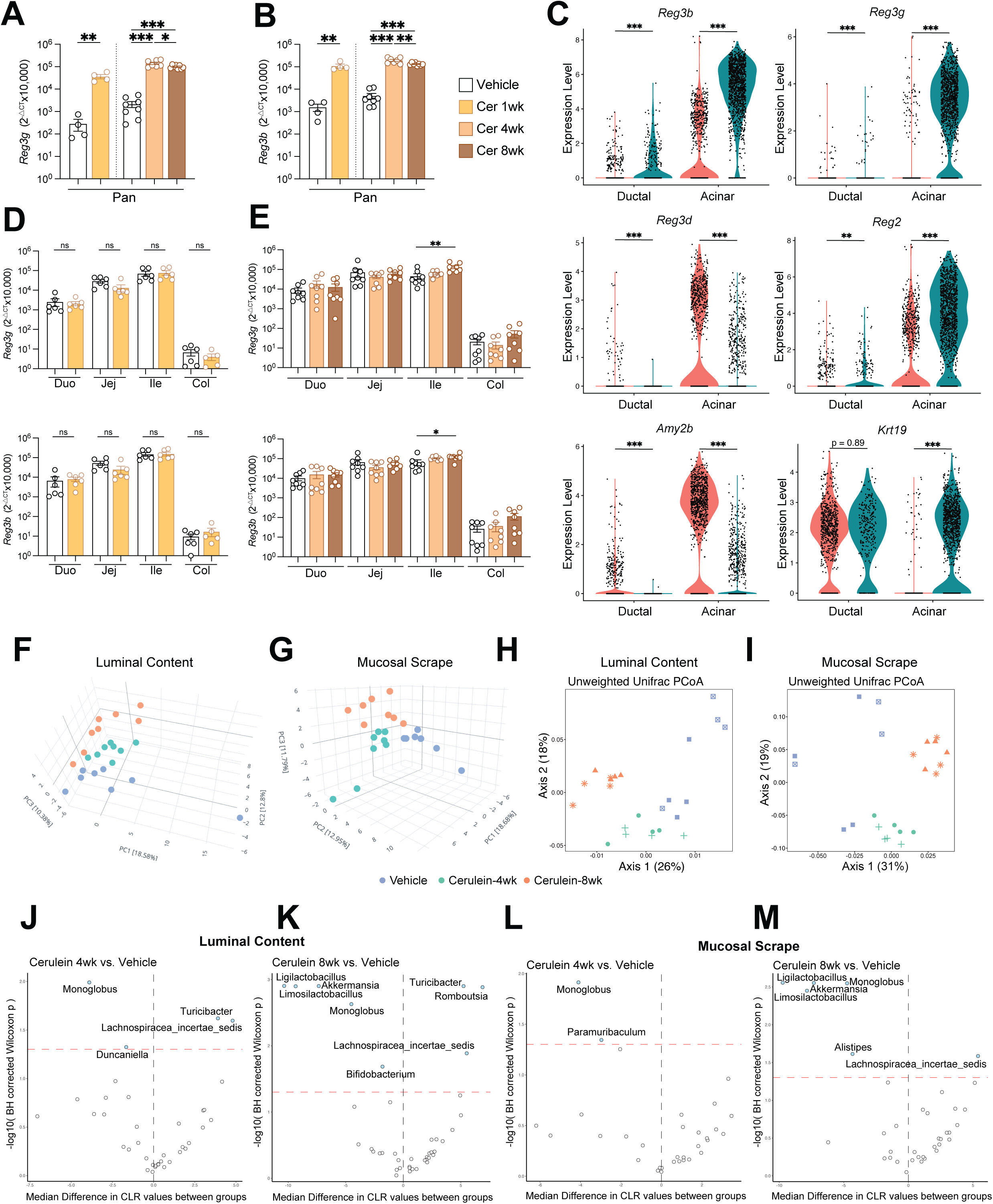
Cerulein induced pancreatitis triggers upregulation of *Reg* genes in pancreas and modifies the gut microbiome. **A, B.** Pancreatic expression of *Reg3g* (**A**) and *Reg3b* (**B**) in mice treated with Cerulein for 1, 4 or 8 weeks and age-matched vehicle control mice housed in parallel (*n*=4 (1 week) or 8 (4 and 8 week treatment)). Data representative of one experiment per time point. **C**. Violin plots of *Reg3b, Reg3g, Reg3d, Reg2, Amy2b* and *Krt19* expressions in mouse acinar versus ductal cells from vehicle or Cerulein treated mice based on *scRNAseq.* **D, E**. Intestinal expressions of *Reg3g* and *Reg3b* in all gut segments from mice treated with Cerulein or vehicle for one week (**D**) or 4 and 8 weeks (**E**) measured by Q-PCR (*n*=6 (1 week) or 8 (4 and 8 weeks)). Data representative of one experiment per time point. **F-I**. PCA of duodenal luminal bacteria (**F**) and mucosa-associated bacteria (**G**) or beta-diversity of duodenal luminal bacteria (**H**) and mucosa-associated bacteria (**I**) of mice with indicated treatments (*n*=7-8). **J-M.** Volcano plots of bacteria enriched or de-enriched in the mucosa of mice treated with Cerulein for 4 (**J**) or 8 (**K**) weeks versus vehicle control mice, or in the lumen of mice treated with Cerulein for 4 (**L**) or 8 (**M**) weeks versus control mice (*n*=8). * p<0.05, ** p<0.01, *** p<0.001 by student t-test.

Despite these dramatic pancreatic changes, the intestinal expression of the two proxy *Reg* isoforms, *Reg3b* and *Reg3g*, remained largely unaltered along the gut (Fig. 5D, E), with only ileal expression increasing 2-fold after 8 weeks. Thus, even in pancreatitis, the signal responsible for increased pancreatic *Reg* expression was not transmitted to the gut. Given the almost immediate effect of the cerulein treatment on *Reg* expression, we wondered if the duodenal microbiome might be affected in this setting. Notably, the fecal microbiome has been described to be altered by cerulein treatment^54^, a finding we confirmed at all time points (Fig. S9A, B, Table 2, 3). When analyzing the microbiota of duodenal luminal contents and mucosal scrapes by 16S rRNA gene sequencing, we also found microbial differences between mock- and cerulein-treated animals by PCA as early as 1 week of cerulein treatment (Fig. 5F, G; Fig. S9C, D, Table 2, Table 3). Beta diversity started to segregate by treatment after 4 and 8 weeks, despite cage effect (Fig. 5H, I; Fig. S9E, F). Total bacterial loads were unchanged (Fig. S9G, H), but the absolute abundance of distinct phyla was altered: After one week of treatment, several gram-positive phyla were enriched in the duodenal mucosa and lumen, including *Turicibacter*, *Lachnospiracea* and *Clostridia* (Fig. S9I, J). This enrichment persisted for *Turicibacter* and *Lachnospiracea* at later time points in the luminal contents, but from 4 weeks onwards, more phyla were depleted than enriched, representing a mix of gram positive and gram-negative bacteria (Fig. 5J-M). While most of these bacteria likely originated from coprophagia when recovered in the duodenum^55^, the results show that in the case of cerulein-induced pancreatitis, the upregulation of *Reg* genes in the pancreas correlates with a change in duodenal microbiome composition.

**Table 2.** Full list of microbial comparison of acute cerulein.

**Table 3.** Full list of microbial comparison of chronic cerulein.

Thus, cerulein-induced pancreatitis, like PDAC, does not induce *Reg* expression in the gut despite strong induction of *Reg* in the pancreas. Though not necessarily mediated by pancreatic REG proteins, the abundance of select bacterial phyla in the duodenum changes in this setting.

## Discussion

In this study, we interrogated the baseline expression profile of all *REG/Reg* isoforms in the pancreas and gut, as well as their regulation upon intestinal and pancreatic perturbations. By analyzing the isoform usage and components of upstream signaling machineries, we uncovered the differential *REG/Reg* isoform usage between pancreas and gut, the similarity between pancreas and duodenum as well as between mice and humans, a common, unique sensitivity to IL-22 in these two organs, and the ability of the gut microbiome and its composition to match intestinal and pancreatic expression of select *Reg* genes. The findings thus suggest a deliberate and coordinated recruitment of pancreatic antimicrobial support by the gut and represent a hitherto underappreciated gut-to-pancreas axis, by which intestinal microbiota regulates pancreatic innate immunity. Furthermore, we classified *Reg* genes into “inducible” and “constitutive” based on their overall regulation by the microbiota and enrichment for STAT3 binding sites. We encountered this classification also in two primary pancreatic diseases. In these settings, we found that *Reg* regulation remained local, possibly because they depended more on IL-6, which is primarily produced by tissue resident immune cells.

### Evolution of divergent REG isoforms and inter-species differences in pancreatic and intestinal usage

In both species investigated here the *REG/Reg* family consists of three subfamilies. The similarity hierarchy aligns with *REG4/Reg4* being the evolutionarily most ancient isoform^23^ that, via gene duplication onto another chromosome, gave rise to the *Reg1* and *Reg3* subfamilies. In both mice and humans, *REG1A/Reg1* dominated the pancreas and duodenum. In mice, the duodenum was also consistently the gut segment most similar to pancreas regarding all other *Reg* isoforms, while *Reg4* and the *Reg3* family, with exception of *Reg3d*, dominated the ileum. This suggest that, if all *Reg* isoforms bind microbial sugars, they are poised to respond to different microbes expected to be encountered in the different gut segments. The pancreas’s similarity to duodenum could serve two purposes: keeping itself sterile from refluxing microbes via the pancreatic duct and supporting duodenal antimicrobial activity through REG1 secretion into the pancreatic juice. The stark difference between baseline *REG1A/Reg1* expression and other isoforms, even in GF mice, raises intriguing questions about what evolutionarily drove this difference. Assuming REG1 indeed has antimicrobial activity, it is possible that a high baseline in the upper intestinal tract guarantees the elimination of microbes at the site where they are least wanted, the gut segment with the highest nutrient levels and absorption. REG1 may be targeting bacteria and fungi that would otherwise outgrow in the comparatively oxygen-rich, low pH environment of the duodenum. The relatively few symbiotic strains inhabiting the duodenum may have evolved resistance to this isoform. By contrast, the isoforms more enriched in the distal intestine would have to be more tightly controlled, as an abundant microbiome is desired in the ileum and colon, yet REG3 isoforms have been shown to target symbiotic bacteria when upregulated ^2,56^. Thus, REG3 may only be upregulated when the benefit of killing microbes prevails, though REG3-insensitive organisms like Salmonella appear to hijack the system and trigger *Reg3* upregulation to clear a niche for themselves^3,57^. Determining the ligand preferences of each isoform, as well as their oligomerization properties, will be crucial to fully appreciate the benefits of differential isoform usage along the gastrointestinal tract, but also how select isoforms could be blocked for therapeutic purposes. Similarly, promoter analysis of the different isoforms and cognate transcription factor levels in pancreas and along the gut warrant further investigation to understand the differences in baseline expression and sensitivity to IL-22.

Given the mouse is the most common model organism for human disease, it is reassuring that in both species *REG1A/Reg1* is the main isoform in the pancreas, total *REG* expression is higher in pancreas than gut with the colon as the lowest site. However, humans differ from mice in their low intestinal baseline expression of *REG3* isoforms, which is reported by others as well. Since newborn and GF mice display similar *REG3* expression patterns, it likely reflects an evolutionary adaptation of the murine gut, possibly driven by their natural coprophagic behavior.

### Conservation of REG/Reg prosegment and its relation to trypsin

Trypsin is a critical activator of REG3A^22^ and likely other REG proteins, as the inhibitory pro-segment and trypsin cleavage site are conserved among all REG proteins. Trypsin, produced as a zymogen with low autocatalytic capacity, requires activation by enterokinase^58^ and is inhibited by trypsin inhibitor proteins^59,60^. This tightly regulated mechanism ensures controlled activation of REG proteins, potentially preventing self-damage that could occur if REG proteins interacted with the mammalian cell surface proteome, which is heavily glycosylated^22^. Of note, viral and bacterial pathogens use lectins to attach to or kill eukaryotic cells^23,25^, and some eukaryotic DAMP receptors^61^ and snake venoms^62^, which bind to eukaryotic surface glycoproteins, are also C-type lectins. Therefore, at high concentrations, REG proteins could damage self-tissues through low-affinity but high-avidity binding to eukaryotic carbohydrates, tethering REG complexes to cell membrane. Trypsin, primarily generated in the pancreas, is expressed at lower levels in the gut epithelium^60^ and could activate REG proteins in gut segments beyond the reach of pancreatic juice. The unequal distribution of trypsin has two significant implications. One, that the difference in REG activity between duodenum versus distal intestine will be even bigger than the transcriptional analysis already predicted (Fig. 1I, J). Trypsin-digesting symbionts in the distal intestine^63^ could further accentuate this difference. The other is that inappropriate activation of trypsin poses a higher risk of REG-mediated self-damage in the pancreas than gut. Indeed, pancreatitis, characterized by tissue trypsin activity and acinar self-damage^64,65^, is a predisposing risk factor for PDAC^50,51^. Increased trypsin activity has also been observed in inflammatory bowel diseases^66^. Both are contexts in which *REG* are upregulated also. Attesting to the fatality of this combination, mice lacking all pancreatic *Reg* isoforms are protected from cerulein-induced pancreatitis^20^. The self-damaging potential of REG proteins could unify the seemingly disparate roles of REG: mitogenic in the pancreas versus antimicrobial in the gut. Plasma membrane disruption elicits a cellular repair response and proliferation^67^, a phenotype that will be more apparent in the pancreas than the highly regenerative gut epithelium, though the mechanism could also contribute to gastrointestinal cancers. Additionally, since membrane disruption goes in hand with Ca^2+^ influx^68^, highly secretory and excitatory cells like pancreatic cells may be affected differently by REG proteins than others. The high trypsin activity associated with pancreatitis could also explain why cerulein-induced pancreatitis impacted the duodenal microbiome more than PDAC, possibly through more active REG proteins.

### Microbe-induced IL-22, REG/Reg, and pancreatic disease

Baseline expression of *Reg* in pancreas and gut is partially independent of IL-22, as observed in GF mice with low IL-22 levels^11,69^. However, microbial induction of REG proteins is largely IL-22 dependent: Many symbionts and pathogens^70,71^ or PAMPs^10^ require IL-22 for their *Reg*-inducing effect in the gut. Our study focused on the role of the symbiotic gut microbiota in regulating pancreatic *Reg* expression. The extrapolation of our findings is that situations that drastically increase intestinal IL-22 production and *Reg* levels, such as certain pathogens, tissue damage or barrier breach, could similarly affect the pancreas, an inference that will be important to address experimentally in the future. Given the link between IL-22 and pancreatic diseases, whether pathobionts or gastrointestinal infections that trigger IL-22 are involved in the initiation or progression of chronic pancreatitis and PDAC in humans is of direct clinical relevance. Primary pancreatic disease was sufficient to increase *REG/Reg*; but the microbiota may set a “permissive tone”. GF and *Il22* knockout mice exhibit drastically slowed PDAC progression^43,72^ while *Reg3g* overexpression is sufficient to accelerate it^19^. Mice lacking gut bacteria are also protected from developing pancreatitis^73^. However, primary dysbiosis, with an overrepresentation of strains efficient at triggering REG upregulation, could conceivably aggravate pancreatic disease progression. In our study, the pancreatic microbiome, which is low in abundance, remained unchanged upon PDAC, as did the duodenal and fecal microbiomes. However, dysbiosis was reported in a stochastic and slower KPC model^49,72^, and fungi or bacteria enriched in the pancreas or feces of PDAC patients can accelerate the disease in the murine model^72,74,75^. Notably, PDAC-aggravating bacteria can elicit IL-17^76^, and many IL-17 secreting cells co-produce IL-22^42,77^, raising the possibility that the same gut bacteria could impact the pancreas via synergistic mechanisms, with IL-17 primarily targeting ductal cells and IL-22 targeting acinar cells, based on our *scRNAseq* analysis, to aggravate PDAC. Regardless of whether live duodenal microbes invade the pancreas, our data based on SFB with strict ileal tropism, suggest that direct invasion is not necessary for the gut microbiota to influence pancreatic innate immunity.

### Lack of reciprocity between pancreas versus gut initiated Reg upregulation

While direct gut-pancreas cooperation is well known in the context of postprandial metabolism, our findings underscore that these organs also coordinate their immune responses^78–81^. In our study, the gut exerted a stronger impact on innate immunity of the pancreas than vice versa. This conceptually aligns with the need for antimicrobials and enzymes in pancreatic juice to support gut defenses. In addition, there is a real threat of ductal microbial reflux from gut to the pancreas. Anatomically and mechanistically, the stronger intestinal influence on the pancreas is logical. The intestine, being a much larger organ, harbors the biggest and densest population of immune cells outside of lymphoid organs, contributes more lymph to shared LNs due to larger lymphatic output, and its venous drainage passes in part through the pancreas. Therefore, whether the coordinated *Reg* transcription is mediated via PAMPs, metabolites, blood-borne cytokines or gut-primed circulating immune cells traveling between organs, the influence is stronger if the gut is the source. Understanding the relative importance of each mentioned pathway in specific situations is imperative for intercepting pathological gut-to-pancreas signals.

### Limitations of this study

Our conclusions are largely based on transcriptional data rather than protein levels. The inferred relative abundances of *REG/Reg* transcripts within organs assume similar efficiency across all Q-PCR primers used. Although all primers were verified and *scRNAseq* data supported our conclusions, there may be slight variations in actual ratios due to subtle differences in primer performance. Our tissue panel reflects *Reg* family and cytokine receptor expression at homeostasis, however, others have reported increased IL-22 signaling upon injury/tissue damage in organs, like liver, skin and lung^70^. Finally, rare cell types within organs that express IL-22 receptors may be below the detection limit in whole tissue extracts.

## Acknowledgements

We would like to thank the University of Chicago Animal Resource Center and microscopy, flow cytometry, functional genomics and Duchossois Family Institute (DFI) microbial sequencing cores and the organ donors to the Gift of Hope. This study was supported by the Pancreatic Cancer Action Network, the Cancer Research Foundation, NIH R01 DK133393, A DDRCC pilot and feasibility grant, the Searle Scholar’s Program, Pew Charitable Trust, University of Chicago start-up funds (DE) and Duchossois Family Institute multidisciplinary grant (DE, MLM). Additional support was provided by NIH T32 AI007090 (MK), a Duchossois Family Institute at University of Chicago postdoctoral fellowship (AF), Jeff Metcalf Fellowships (FS, GL and ON), the UChicago Careers in Healthcare Katen Scholars program (FS), the Chicago EYES on Cancer program (KM), the UChicago Quad Undergraduate Research Scholars Program (ON), a National Science Foundation Graduate Research Fellowship, DGE-1745301 (NJW-W), a Caltech John Stauffer SURF Fellowship (PN), and a grant from Caltech’s Beckman Foundation (RFI).

## Author contributions

DZ-conceptualization, most experiments, figures; MK-Taconic and Jax mice co-housing experiment, immunofluorescent staining and scRNAseq analyses; FS-multi-tissue analysis and additional RNA extractions; GL, KM and ON-RNA extractions and Q-PCR; ENO-Il22 knockout mouse experiment; NJWW-contributed in conceptualization of microbiota study with DZ/RFI, NJWW and PMN-16S rRNA gene sequencing and analysis for duodenal samples for 12 week PDAC and 1week Cerulein/vehicle samples (shown in figures S7J-K & S9C-F, I-J); AF-human intestine collection and and scRNAseq analysis; JK, NC, and MLM-human intestine collection, PW-human pancreas collection, RFI-contributed to conceptualization of microbiota study, DE-project oversight, experiments, funding, first draft. All authors-read and edited.

## Materials and Methods

### Reagents

The following reagents were used: Trizol (Thermo Fisher Scientific), Qubit RNA IQ Assay Kit (Thermo Fisher Scientific), Superscript IV (Thermo Fisher Scientific), Cerulein (Bachem), Trichome (Sigma), Sheep anti-mouse REG3B (R&D, AF5110), Rat anti-mouse CK19 (DHSB, AB2133570), Goat anti-mouse CPA1 (R&D, AF2765), AF488 donkey anti-sheep (Jackson Immunoresearch, 713-545-003), Cy3 donkey anti-rat (Jackson Immunoresearch, 712-165-153), AF647 donkey anti-goat (Jackson Immunoresearch, 705-605-147), DAPI Fluoromount G Clear Mounting Media (SouthernBiotech).

### Human specimens

Human pancreas samples were obtained through the Gift of Hope organ donor program (Dr. Piotr Witkowski). Human gut samples were obtained through the Gift of Hope organ donor program (Drs. Jonathan Kent and Lucia Madariaga). The studies were deemed exempted from further IRB review by the University of Chicago BSD IRB on the basis of them being secondary research on material from deceased and de-identified subjects (IRB19-1942 and IRB19-1085).

### Mice

All experiments were performed in accordance with the University of Chicago ACUP. C57BL/6J, NOD/ShiLtJ, Ptf1a^CreERT2/w^, *Trp53*^fl/fl^, and Kras^LSL-G12D^, *Il22*-/- were purchased from The Jackson Laboratory and crossed and maintained at the University of Chicago animal facility adhering to the IACUC protocol under specific pathogen-free (SPF) conditions, with the additional stipulations of the mice being free of segmented filamentous bacteria (SFB), murine norovirus (MNV) and Hpp. Germ free (GF) mice were purchased from Taconic and maintained in isolators in the gnotobiotic facility at the University of Chicago. Ex-GF mice in our study describe the offspring of GF mice that had been taken out of our isolators, colonized with microbiota from Jax mice mated under our strict SPF conditions for at least two generations with the same bedding and food as provided in isolators.

### Pancreatic cancer induction

Ptf1a^CreERT2/w^*Trp53*^fl/fl^Kras^LSL-G12D/w^ (KPC) mice and their wildtype littermates (Ptf1a^w/w^*Trp53*^fl/fl^Kras^LSL-G12D/w^ or Ptf1a^w/w^*Trp53*^fl/fl^Kras^w/w^) were fed with tamoxifen-containing diet (Envigo # TD.130858) for 7 days at 3-4 weeks of age for cancer induction. Mice were sacrificed at indicated time points. Pancreas and different intestinal segments were taken for subsequent processing and analysis.

### Model of acute and chronic cerulein-induced pancreatitis

The acute pancreatitis was induced by 6 hourly intraperitoneal injections of 50ug/kg of the CCK analogue cerulein on each injection day, 3 injection days per week for 1 week. The chronic pancreatitis was induced by 6 hourly intraperitoneal injections of 50ug/kg cerulein on each injection day, 2 injection days per week for 4-8 weeks. Control mice were injected with the same amount of saline.

### REG protein sequence alignment

Protein sequences of mouse and human REG family members were obtained from the UniProt database (uniprot.org). Multiple sequence alignment was performed using the built-in alignment tool in UniProt, with the number of iterations set to 3. Sequence similarity matrix was obtained from the alignment output.

### REG protein structural prediction and RMSD calculation

Trypsin cleavage sites were identified in the amino acid sequences of each REG protein. The resulting truncated sequences were input into the “Monomer structure prediction with AlphaFold” module on Tamarind Bio (tamarind.bio/alphafold). To test for module accuracy, the predicted structures were superimposed onto the predicted full-length structures extracted from AlphaFold database (alphafold.ebi.ac.uk). Root mean square deviation (RMSD) of atomic positions was calculated between each REG protein using PyMOL (Version 2.5.8).

### Cell culture

Mammalian cells were all grown in a humidified incubator with 5% CO_2_ at 37°C. PANC1 cells (ATCC) were grown in Dulbecco’s Modified Eagle Media (DMEM, Gibco) supplemented with 10% FBS and 100 U/mL penicillin and streptomycin. ASPC1 cells (ATCC) were grown in RPMI-1640 media supplemented with 10% FBS and 100 U/mL penicillin and streptomycin.

### Tissue procurement, RNA extraction and Real-time PCR

Mouse pancreatic and intestinal tissues were frozen in RNAlater and stored at −80 °C. Human pancreata were stored as entire organs at −20C for less than 12 months. Human intestinal pieces were frozen in RNAlater and stored at −80 °C, and only the mucosa processed for RNA extraction. Total RNA from samples was extracted using TRIzol, RNA integrity was verified by Qubit RNA IQ Assay and cDNA generated by reverse transcription using SuperScript IV kit according to the manufacturer’s protocol. Quantitative real-time PCR was performed using Power SYBR Green PCR Master Mix (Applied Biosystems) on Quanstudio 6 Flex machine (Thermo Fisher Scientific). The relative expression of target genes was normalized to *36B4* (mouse) or *B-ACTIN* (human) expression and calculated using the formula of 2^−ΔCt^ x10000. All data were averaged from at least 2 replicates. Newly designed primers were verified for single peak melting curves, single band products on an agarose gel and linearity within the range of measured CT values. The primers used were:

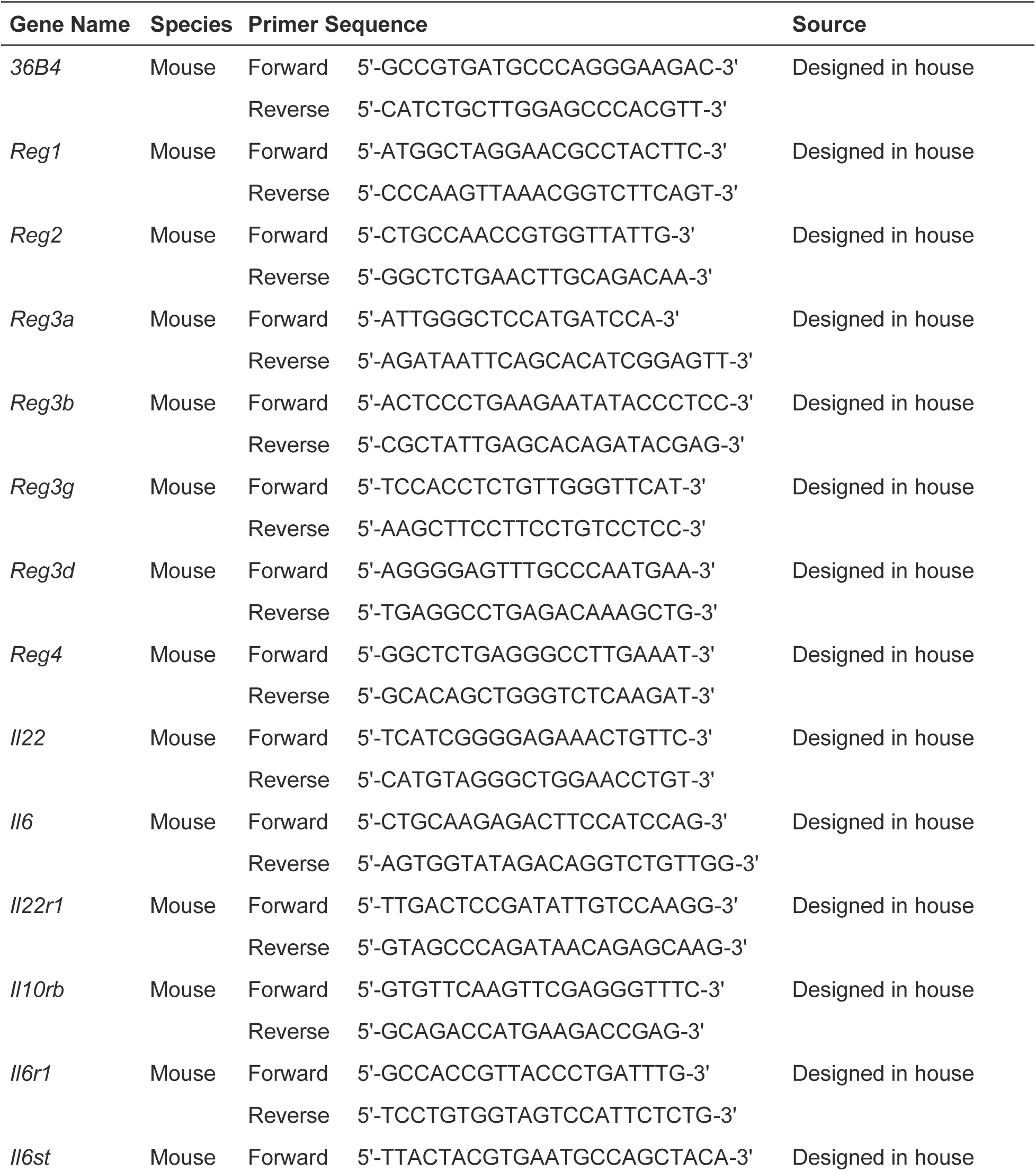
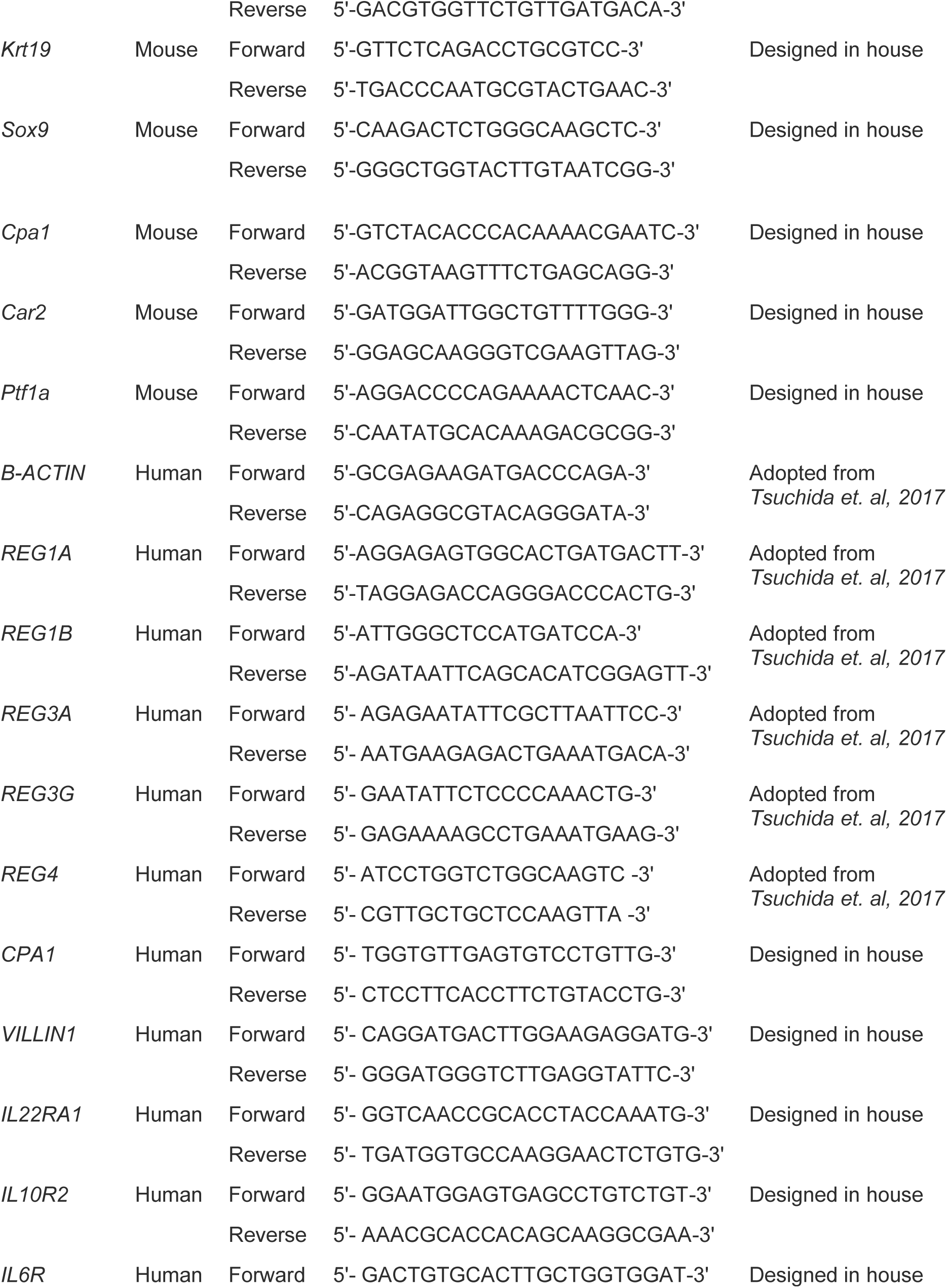
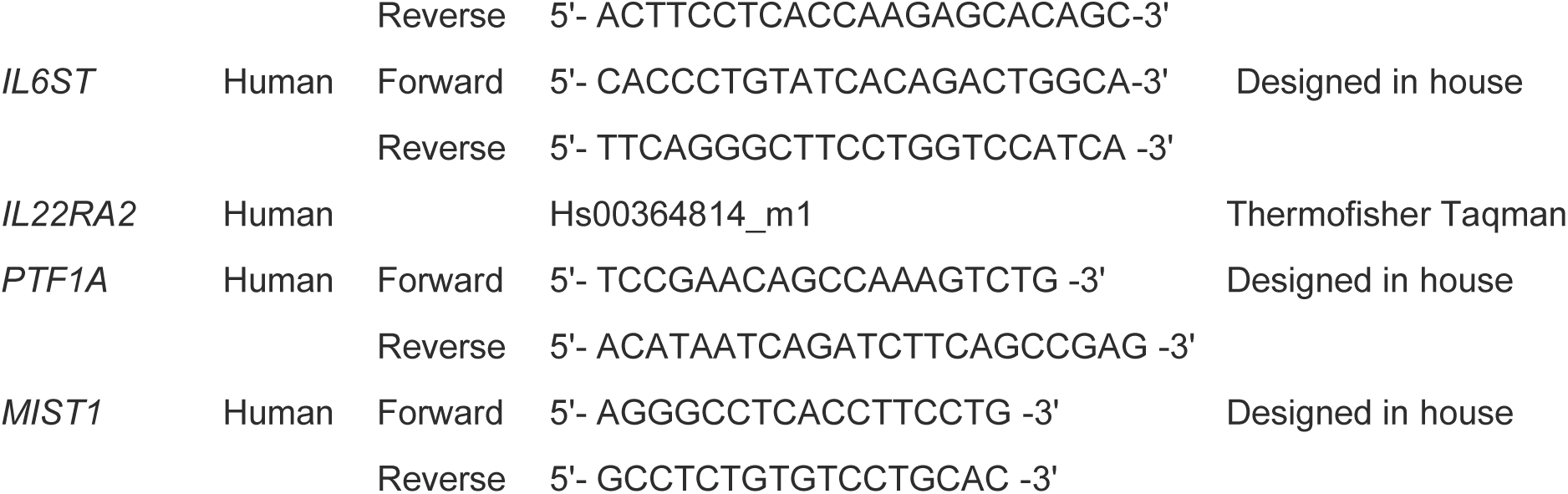

### Histological imaging and assessment

Specimens were immediately collected from mice and fixed in 4% paraformaldehyde for 1 hour at room temperature and rinsed in 70% ethanol. All the samples were graded-ethanol dehydrated, embedded in paraffin, cut into 5-μm–thick sections using microtome and then stained with hematoxylin and eosin (H&E) or special Masson trichrome for histomorphometric assessment at University of Chicago Medicine Human Tissue Resource Center. The stained sections were sent to University of Chicago Microscopy Core for whole slide scanning using Olympus VS200 slideview research slide scanner at 20x resolution.

### Immunofluorescent staining

5-μm-thick paraffin-sections were heated to 65°C for 20min and rinse twice in xylene for 10min to de-paraffinize. Slides were re-hydrated using graded-ethanol and then blocked for 30 min at RT with blocking buffer (5% donkey serum, 2.5% BSA in 1X PBS). Slides were permeabilized with 0.1% Saponin if necessary, then incubated overnight in a humidified chamber at 4°C with antibodies (REG3β and CPA1 at 1:100, CK19 at 1:50 dilution). The next day, slides underwent three PBST (1X PBS with 0.05% Tween-20) washes, and were incubated with appropriate secondary antibodies (1:500 dilution) for 1hr at RT. After three more PBST washes, slides were mounted using DAPI Fluoromount G Clear Mounting Media. Stained slides were sent for whole slide scanning.

### Bioinformatic analysis of published scRNAseq datasets

scRNAseq data was obtained from the NCBI GEO database. Counts were used as input for the R package Seurat. Cells were initially filtered to remove doublets and poor-quality cells based on the unique molecular identifiers (UMIs), number of features, and percentage of mitochondrial genes (<0.25). To normalize gene counts, Seurat’s LogNormalize function was used with default parameters (scale factor = 10,000). Variable genes were then identified using the Seurat function FindVariableFeatures with default parameters. The data was scaled using ScaleData and then subjected to dimensionality reduction by PCA using RunPCA. The cells were clustered using Seurat’s FindNeighbors followed by FindClusters prior to visualization by RunUMAP. For mouse wildtype and cerulein-treated mouse pancreatic scRNAseq dataset GSE188819^32^, clusters annotations were included in the counts file downloaded from GEO database. For mouse pancreatic cancer scRNAseq data GSE141017^82^, logFold changes of genes of interest between acinar and metaplastic cells were extracted and plotted. For human scRNAseq data from GSE155698^33^, clusters were annotated based on the gene set presented in Figure 2b of Steele et al, 2020^33^. For the plots shown in Figure S7A, data integration was performed using IntegrateLayers function in Seurat. Integrated data was then downsampled to visualize the relative contribution of each cell type. For intestinal epithelial cell scRNAseq data from GSE185224 (human)^35^ and GSE201859 (mouse)^34^, processed Seurat objects were downloaded from https://cellxgene.cziscience.com/datasets. Specific expression was visualized using FeaturePlot and VlnPlot functions. Statistical analysis was performed using the ggpubr package in R by t-test.

### Collection of duodenal content and mucosal scrape

The small intestine was removed and the duodenum segment (the first quarter) was identified. Luminal contents were collected by gently squeezing the duodenum. The duodenum was then opened longitudinally and the mucosal surface was scraped using a curled forcep. Both fractions were collected into sterile eppendorf tubes and weighed. The tools were cleaned after each collection.

### Duodenal host depletion, extraction, library preparation, sequencing, and analysis

The host depletion method was adopted from *Wu-Woods et. al, 2023*^83^. In summary, duodenal luminal contents and mucosal scrapes were placed into 2-ml 1.4-mm ceramic bead-beating tubes (Lysing Matrix D from MP Biomedical, catalog no. 1169130-CF) and 400 μL of sterile 0.9% saline was added into the bead-beating tube. Samples were homogenized for 30s at 4.5 m s^−1^ on Qiagen’s TissueLyser II and then immediately placed on ice. 150 μL of homogenized tissue was placed into clean microfuge tubes containing 10 μL of buffer (100 mM Tris + 40 mM MgCl_2_, pH 8.0 and 0.22 μm sterile filtered), 33 μL of saline (0.9% NaCl, autoclaved), 2 μL of Benzonase Nuclease HC (EMD Millipore catalog no. 71205) and 5 μL of Proteinase K (NEB catalog no. P8107S). Tubes were incubated for 15 min at 37 °C with shaking at 600 rpm, then pelleted at 10,000g for 2 min. The supernatant was discarded and pellets were resuspended in 150 μL of PrimeStore MTM (Longhorn) to inactivate residual enzymatic activity and stored at −80 °C until shipment to Caltech.

Nucleic-acid extraction was performed using Qiagen’s AllPrep PowerViral DNA/RNA Kit (catalog no. 28000-50) using 3 rounds of bead beating for 1 min at 6.0 m s^−1^ per round on the FastPrep-24 MP Bio Homogenizer. Reagent DX (0.5% v/v; Qiagen catalog no. 19088) was added to PM1 prior to bead beating to reduce foaming.

16S rRNA gene Quantitative Sequencing (Quant-Seq) was performed as previously described^84^. In summary, the V4 (515F-806R) region of the 16S rRNA gene was amplified in duplicate using 5μL template in a total reaction volume of 20μL. Libraries were amplified to ∼15,000 RFU before removal and duplicate reactions were pooled. Sequencing was performed by Fulgent Genetics using the Illumina MiSeq, v2 Standard, 2×250bp.

For microbial abundance measurements, qPCR and ddPCR were performed using the same V4 (515F-806R) primer set. For ddPCR, as previously described^84^, the QX200 droplet dPCR system (Bio-Rad catalog no. 1864001, 1864002) was utilized with 2.5μL of template in a total reaction volume of 25 μL. For qPCR, 1uL of template was added to the following components (final concentration reported): 1X SSOFast EvaGreen Supermix (Bio-Rad catalog no. 1725201), 500 nM forward and reverse primer (F: 5’- CAG CMG CCG CGG TAA -3’, R: 5’- GGA CTA CHV GGG TWT CTA AT -3’) for a total reaction volume of 10 μL. Amplification and quantification was performed on the CFX96 RT-PCR machine (Bio-Rad) under the following cycling conditions: 94 °C for 3 min, up to 40 cycles of 94 °C for 45 s, 52 °C for 30 s, and 68 °C for 30 s. Conversion from Cq to copies per μL was performed using a standard curve and validated with ddPCR. Sequencing analysis was performed as previously described^84^. In brief, processing was performed with QIIME2 (v2022.8) and taxonomy assigned with SILVA 138 99% OTU 515F-806R reference database using unrarefied data. Dada2 trimming parameters were set to the base pair where average quality scores dropped below 30. Raw sequence data, ASV abundance and taxonomy tables, were generated. All downstream analysis was performed at the genus-level. Taxa below LOD (limit of detection) were identified based on Poisson statistics, as previously described^85^, and set to zero in downstream analysis performed in Python (v3.9.12).^83^

### Fecal DNA extraction, 16S library preparation, sequencing and analysis

Fecal microbial DNA was extracted using the QIAamp PowerFecal Pro DNA kit (Qiagen). Prior to extraction, samples were subjected to mechanical disruption using a bead beating method using Qiagen’s TissueLyser II. Samples were then centrifuged, and supernatant was resuspended in CD2, that effectively removes inhibitors by precipitating non-DNA organic and inorganic materials including polysaccharides, cell debris and proteins. DNA was purified routinely using a silica spin column filter membrane. By adding solution CD3, a high-concentration salt solution, DNA selectively binds to the membrane which is recovered using elution buffer. DNA is then quantified using Qubit. 16S Q-PCR was used to determine the total bacterial load in each sample.

The V4-V5 region of 16S rRNA gene was amplified using universal bacterial primers – 563F (5’- nnnnnnnn-NNNNNNNNNNNN-AYTGGGYDTAAA-GNG-3’) and 926R (5’-nnnnnnnn-NNNNNNNNNNNN-CCGTCAATTYHT-TTRAGT-3’), where ‘N’ represents the barcodes, ‘n’ are additional nucleotides added to offset primer sequencing. The approximately ∼360 bp amplicons were then purified using bead-based size selection, quantified, and pooled at equimolar concentrations. Illumina sequencing-compatible Combinatorial Dual Index (CDI) adapters were ligated onto the pools using the QIAseq 1-step amplicon library kit (Qiagen). Library QC was performed using Qubit and Tapestation and sequenced on Illumina MiSeq platform to generate 2×250bp paired-end reads.

Dada2 (v1.18.0) was used as the default pipeline for processing MiSeq 16S rRNA reads with minor modifications in R (v4.0.3). Specifically, reads were first trimmed at 190 bp for both forward and reverse reads to remove low-quality nucleotides. Chimeras were detected and removed using the default consensus method in the dada2 pipeline. Then, ASVs with lengths between 320 bp and 365 bp were kept and deemed as high quality ASVs. Taxonomy of the resultant ASVs was assigned to the genus level using the RDP classifier (v2.13) and trainset 18 (release 11.5) with a minimum bootstrap confidence score of 80. Raw sequence data, ASV abundance and Taxonomy tables, and bar plots, were generated. Samples below 1000 reads and features present in 2 or fewer samples were excluded from the analysis. The Kruskal-Wallis test was used to check significance. Volcano plots were generated using results from the R package ALDEx2. Alpha and beta diversities were estimated using Phyloseq on the repeatedly rarefied datasets (100 iterations) using the R package mirlyn.

### Statistical analysis Q-PCR data

Statistical analyses were performed using *GraphPad Prism* software (version 10.0; GraphPad, La Jolla, Calif). Graphs were generated using *GraphPad Prism* or *Rstudio*. Data are displayed as mean (SD) for at least 3 replicates. Comparison between 2 groups was performed using Student *t* test, whereas comparison among multiple groups was performed using 1-way analysis of variance. A *P* value < 0.05 indicates statistical significance.

---

**Figure S1.**
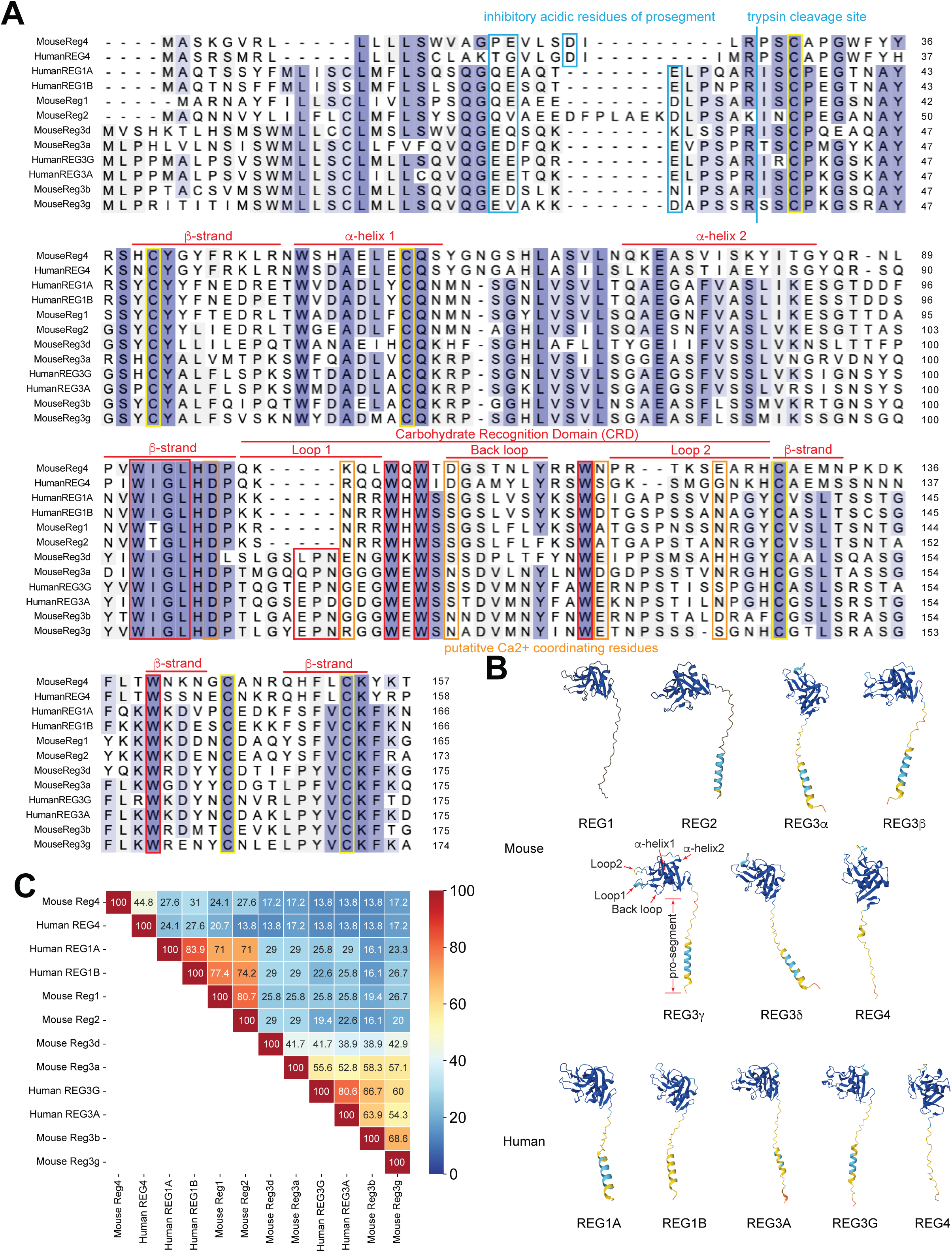
Mouse *Reg* and human *REG* gene family members are structurally conserved. **A**. Amino acid sequence alignment of all mouse and human *Reg* gene family isoforms with the following characteristics annotated: position of acidic amino acid residues critical for inhibitory action of prosegment (one acidic residue sufficient for this property), and prosegment trypsin cleavage site (blue boxes and line), six cysteine residues for protein stabilization via disulfide bridges (yellow boxes), putative D/N/E residues used for coordinating Ca^2+^ ions for canonical ligand binding (orange boxes), features of the C-type lectin fold, notably two predicted alpha helices, five beta strands, a carbohydrate recognition domain (CRD) enriched in tryptophan (W) residues and forming a two-outward (Loop1 and Loop 2) one back-loop structure preceded by the WIGL motif, and the mannose binding EPN motif (red). **B**. Predicted structures of all mouse and human REG proteins (Alphafold), with key features of all REG proteins pointed out on REG3g. **C**. Sequence similarity between the carbohydrate recognition domains of all mouse and human REG proteins.

**Figure S2.**
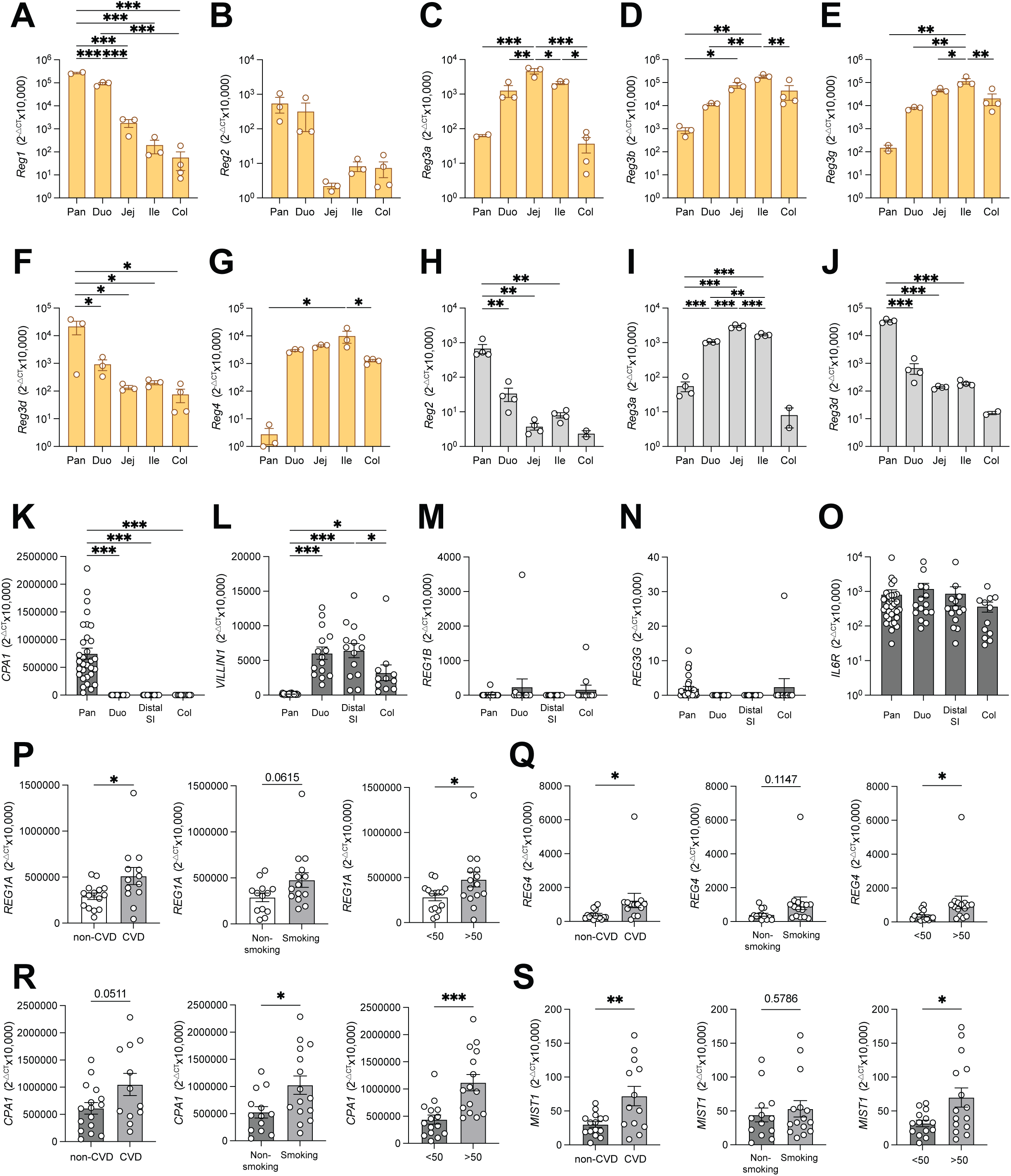
Mouse Reg and human *REG* gene family members differ in their expression patterns between pancreas and gut. **A-G**. Expression levels of indicated *Reg* genes in pancreas (pan), duodenum (duo), jejunum (jej), ileum (ile) and colon (col) of NOD mice by Q-PCR (*n*=3 for all but col, where *n*=4). **H-J**. Expression levels of indicated *Reg* genes in pancreas (pan), duodenum (duo), jejunum (jej), ileum (ile) and colon (col) of C57Bl6 mice by Q-PCR (*n*=4 for all but col, where *n*=2). **K-N**. Expression of *CPA1* (**K**), *VILLIN1* (**L**), *REG1B* (**M**) and *REG3G* (**N**) in human pancreas (pan, n=31), duodenum (duo, *n*=15), distal small intestine (distal SI, *n*=14), and colon (col, *n*=11) by Q-PCR. **O-Q**. Expression of *REG1A* in pancreas of donors with cardiovascular disease (CVD) or not (**O**), smoking history or not (**P**), and below vs above age of 50 at age of death (**Q**). **R-T**. Pearson correlations between expression of indicated genes within same patients. **U-W**. Principal component analysis based on all *REG* genes and *CPA1* in human pancreas, color coded by CVD state (**U**), smoking history (**V**) and age (**W**). * p<0.05, ** p<0.01, *** p<0.001 by ANOVA for multivariant plots (A-N), * p<0.05, ** p<0.01, *** p<0.001 by 2-tailed t-test for two-way comparisons (O-Q).

**Figure S3.**
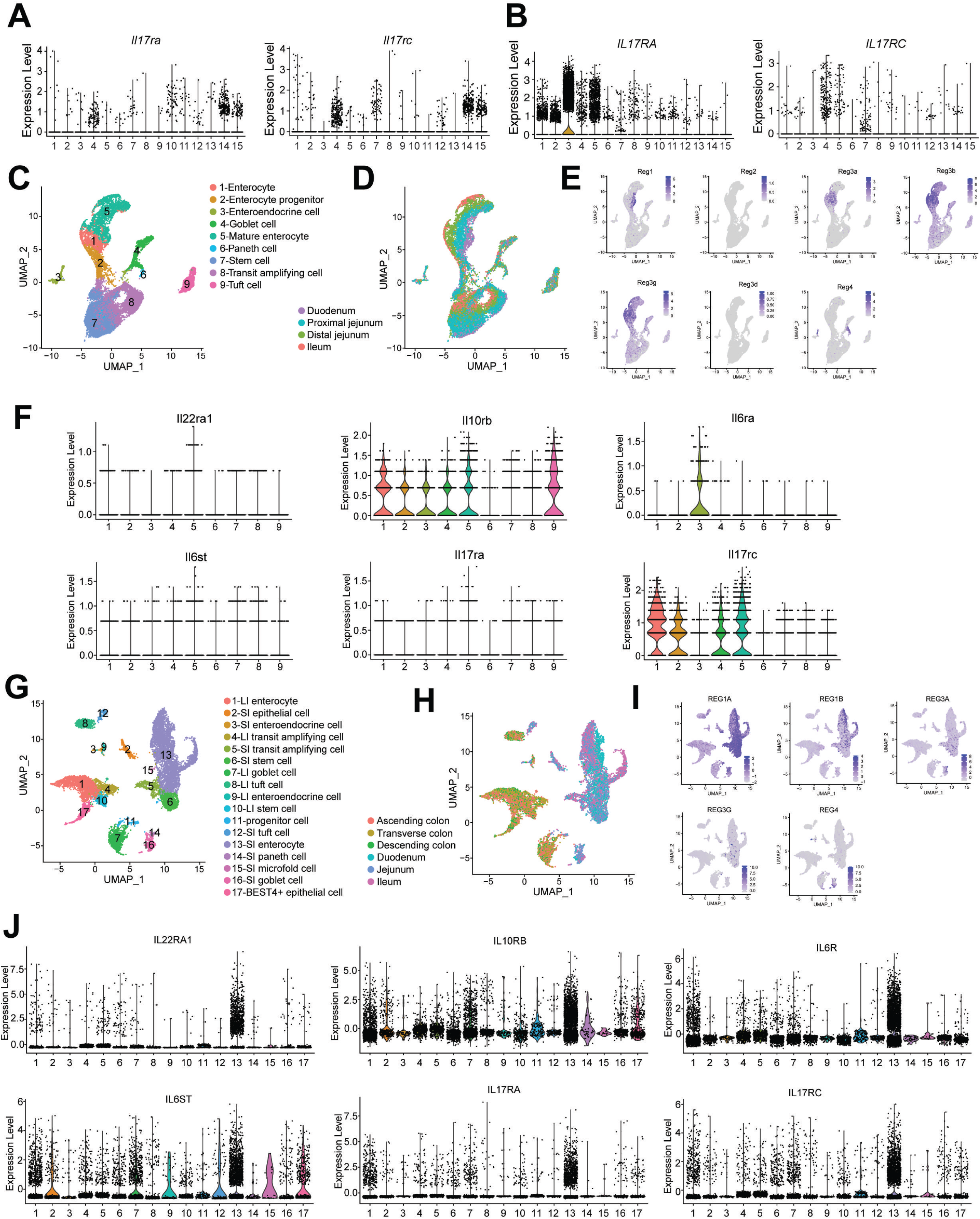
Gut and pancreas are enriched for *Reg* transcripts and the upstream signaling receptors. **A, B.** Violin plots of IL-17 receptor genes across mouse pancreatic cell types (**A**) and human pancreatic cell types (**B**) based on *scRNAseq*. **C, D**. Annotated *scRNAseq* feature plots of mouse small intestine color coded by cell type (**C**) or gut segment of origin (**D**). **E**. Feature plots of mouse *Reg1, Reg2, Reg3a, Reg3b, Reg3g, Reg3d* and *Reg4.* **F**. Violin plots of IL-6, IL-22 or IL-17 receptor subunit genes across mouse small intestinal cell types. **G, H**. Annotated *scRNAseq* feature plots of human intestine color coded by cell type (**G**) or gut segment of origin (**H**). **I**. Feature plots of human *REG1A, REG1B, REG3A, REG3B* and *REG4.* **J**. Violin plots of IL-6, IL-22 or IL-17 receptor subunit genes across human intestinal cell types.

**Figure S4.**
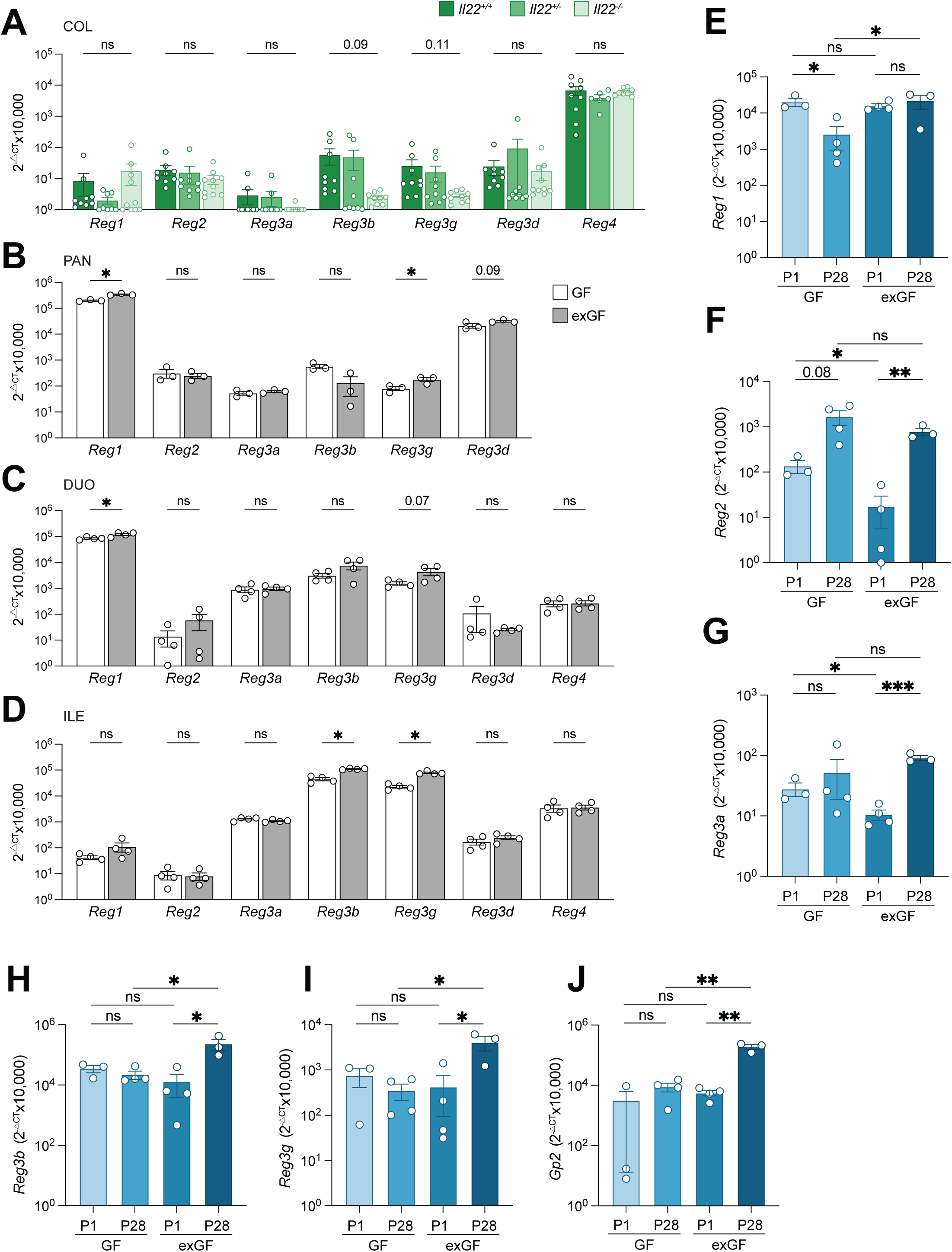
Dependence of baseline pancreatic and intestinal Reg expression on IL-22 and postnatal environmental exposure. **A-D.** Expression levels of indicated *Reg* genes in pancreas, duodenum and colon of *Il22*^+/+^, *Il22*^+/-^ and *Il22*^−/−^ mice (*n*=9, 9 and 10, respectively) by Q-PCR. Data pooled from two independent experiments. **E-J**. Pancreatic expressions of indicated antimicrobial genes in germfree (GF) or housing matched specific pathogen free (exGF) C57Bl6 mice harvested at P1 or P28 by Q-PCR (*n*=3-4, as indicated by white circle dots). * p<0.05, ** p<0.01, *** p<0.001 by student t-test.

**Figure S5.**
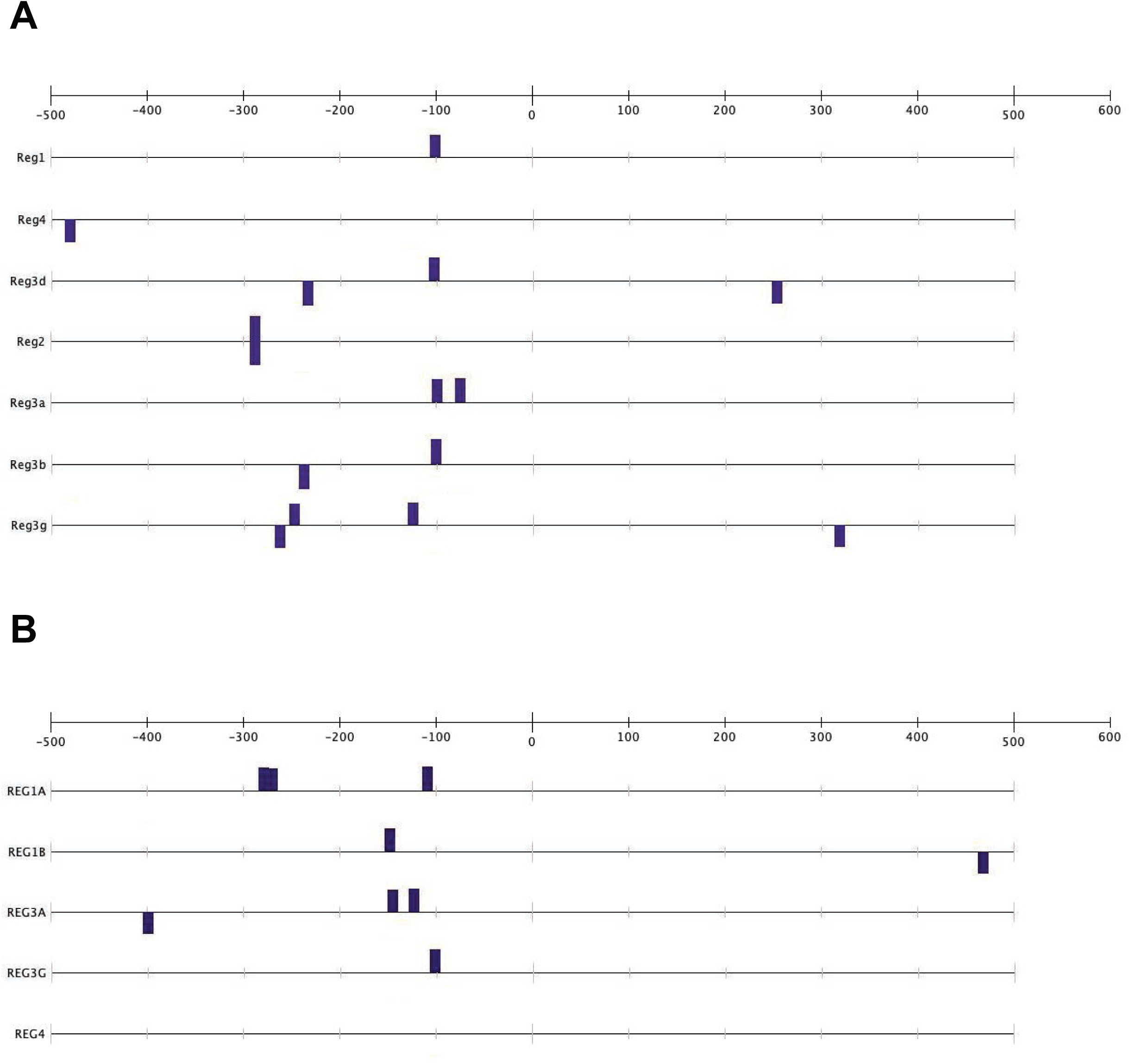
Transcription binding site analysis of REG/Reg family members. **A-B.** Binding site analysis of STAT3 at the promoter region of indicated mouse *Reg* (**A**) and human *REG* (**B**) genes.

**Figure S6.**
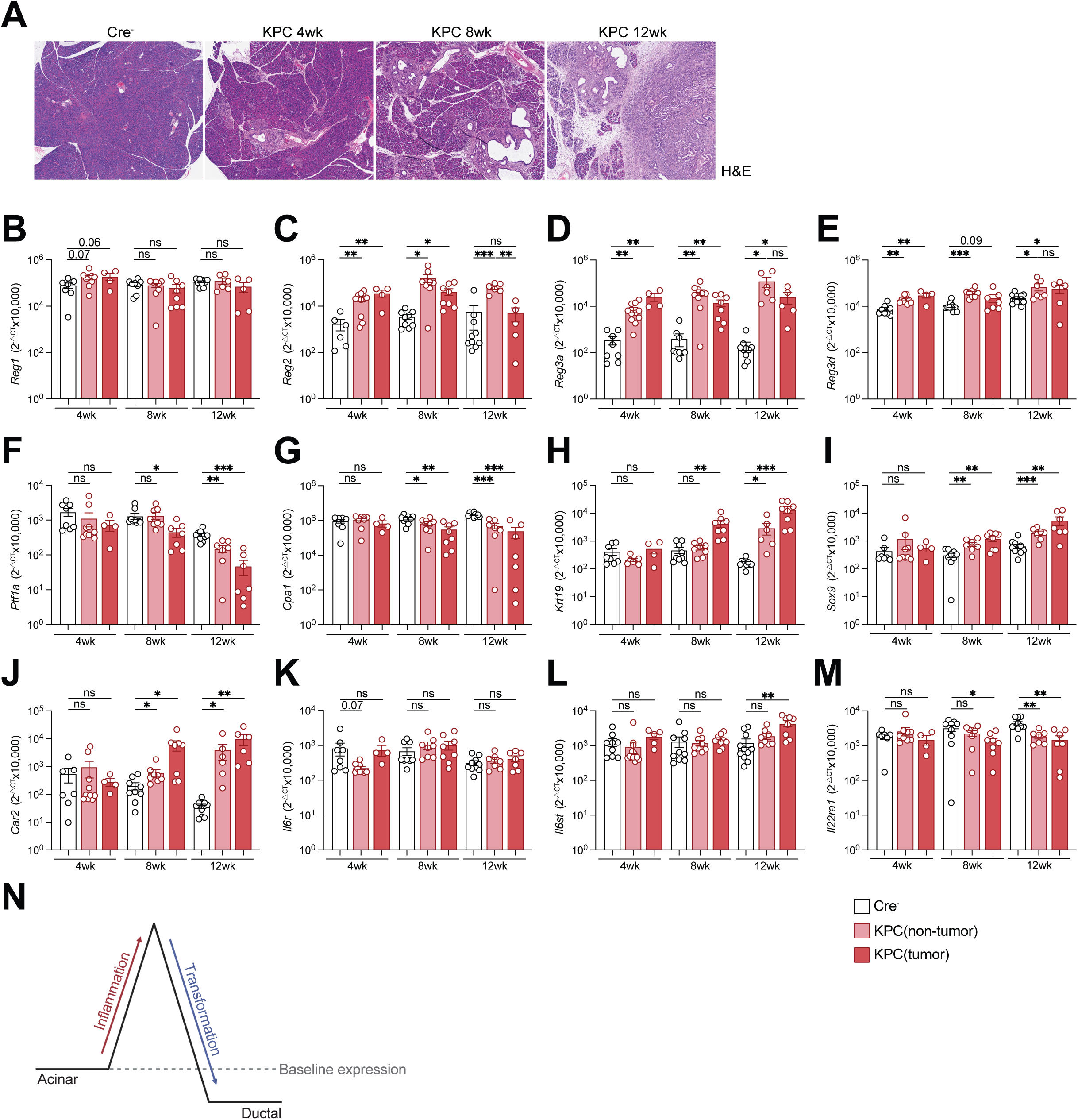
A mouse model of primary pancreatic ductal adenocarcinoma triggers *Reg* upregulation in pancreas but not gut. **A.** Representative H&E-stained paraffin sections of pancreas from KPC mice at 4-, 8- or 12-weeks post tamoxifen treatment and Cre^−^ mice (littermate to 12-week KPC mice). **B-M.** Pancreatic expression of *Reg1*(**B**), *Reg2* (**C**), *Reg3a* (**D**), *Reg3d* (**E**), *Ptf1a* (**F**), *Cpa1*(**G**), *Krt19* (**H**), *Sox9* (**I**), *Car2*(**J**), *Il6r* (**K**), *Il6st* (**L**) and *Il22r1a* (**M**) in KPC mice or Cre^−^ littermates at 4-, 8- or 12-weeks post tamoxifen-induced PDAC onset, measured by Q-PCR (WT *n*=7, 9, 10, PDAC *n*=9, 8, 6 at 4, 8, 12 weeks, respectively). Data pooled from 3 cohorts per time point. **N**. Proposed *REG/Reg* expression during the course of PDAC progression. * p<0.05, ** p<0.01, *** p<0.001 by student t-test.

**Figure S7.**
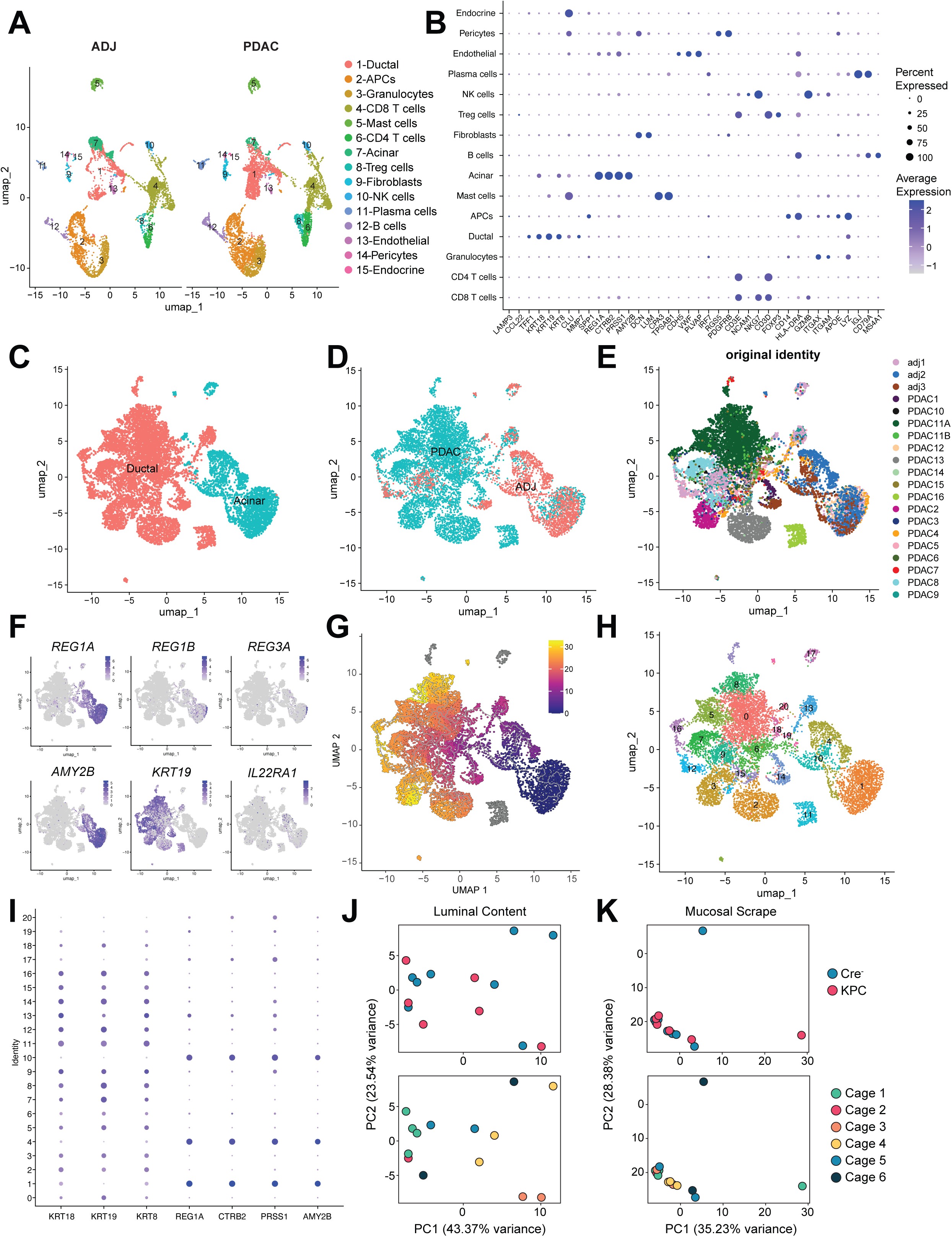
Human PDAC scRNAseq analysis for *REG* genes and murine duodenal microbiome upon PDAC induction. **A, B.** Downsampled UMAPs of cell type clusters (**A**) and dot plot of cluster defining genes (**B**) from pancreatic cells isolated from human PDAC or adjacent tissues analyzed by *scRNAseq*. **C-I.** UMAPs of only acinar and ductal cells coded by cell type (**C**), sample type (**D**), donor sample (**E**), indicated genes (**F**), pseudotime analysis (**G**) and cluster number (**H**) of cells from PDAC and adjacent tissues. **I**. Dot plot of *REG* genes and acinar/ductal marker gene expressions in each cluster identified in **H**. **J**, **K**. PCA of CLR (centered log-ratio transformation) of relative abundance duodenal Luminal Content (**J**) and Mucosal Scrape (**K**) from KPC (*n*=6) or Cre^−^ littermates (*n*=8) taken 12 weeks post tumor induction. Each PCA is coded once by genotype (top) and once by cage identity (bottom).

**Figure S8.**
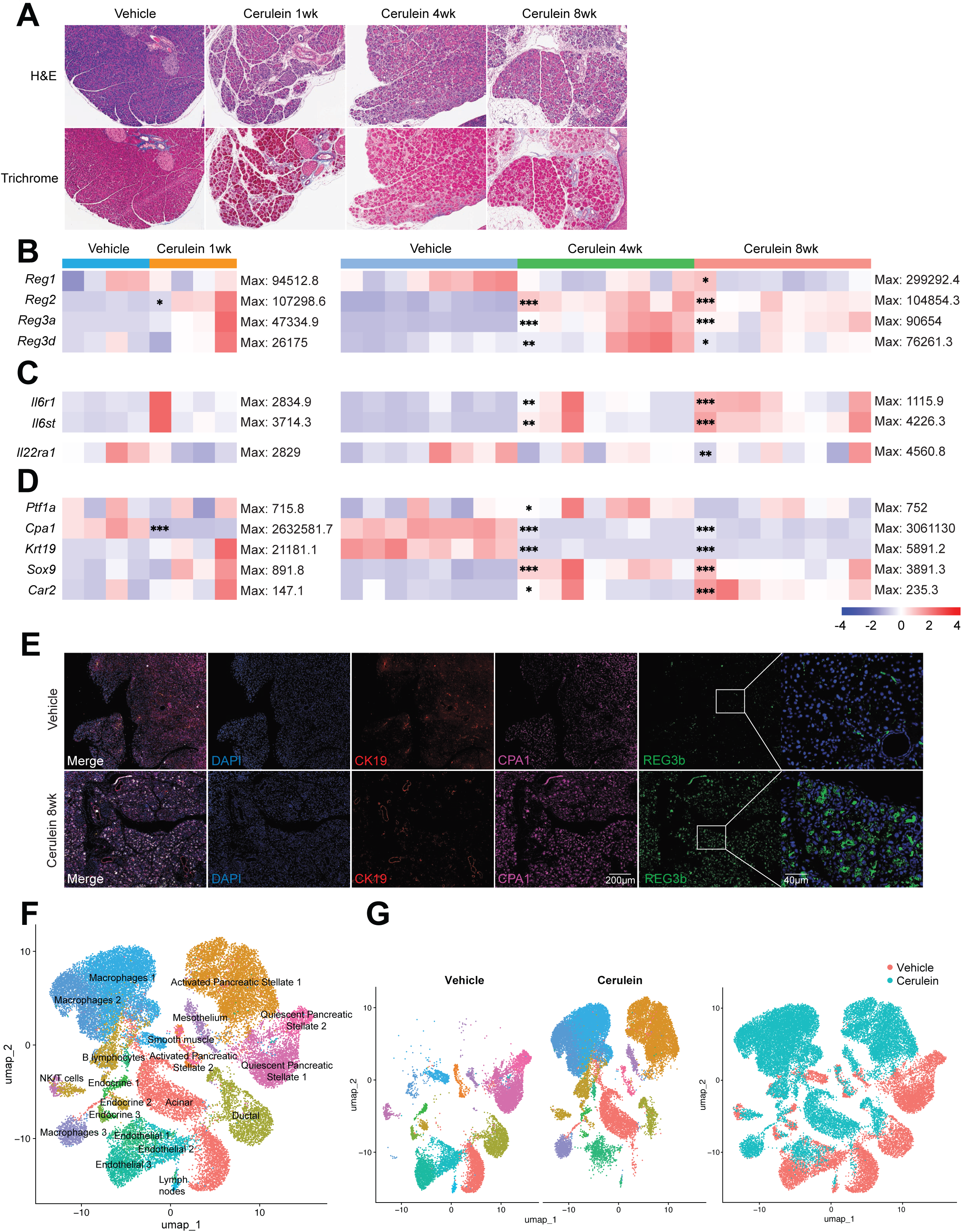
Cerulein induced pancreatitis triggers upregulation of *Reg* genes in pancreas but not gut. **A.** Representative H+E- (top) and trichome- (bottom) stained paraffin sections of pancreata from mice with indicated treatments. **B-D.** Heatmap of expressions of indicated *Reg* genes (**B**), interleukin receptor genes (**C**) and acinar/ductal marker genes (**D**) in pancreata from mice treated with cerulein for 1, 4 or 8 weeks and age-matched vehicle treated mice measured by Q-PCR (*n*=4 (1 week) or 8 (4 and 8 weeks)). Data representative of one experiment per time point. Maximal expression level per row indicated at the right of each row. **E.** Pancreatic sections from mice treated with cerulein or vehicle for 8 weeks stained for ductal marker CK19, acinar marker CPA1, REG3β and DAPI (nuclei). Bar=200 µm, inset bar=40 µm. **F**, **G.** UMAPs of cell type clusters (**F**) and downsampled (left) and color coded by contribution by cells (right) (**G**) from pancreata of vehicle- or cerulein-treated mice based on *scRNAseq*. * p<0.05, ** p<0.01, *** p<0.001 by ANOVA (4 and 8 weeks cerulein comparisons) or two tailed t-test (1 week cerulein comparisons).

**Figure S9.**
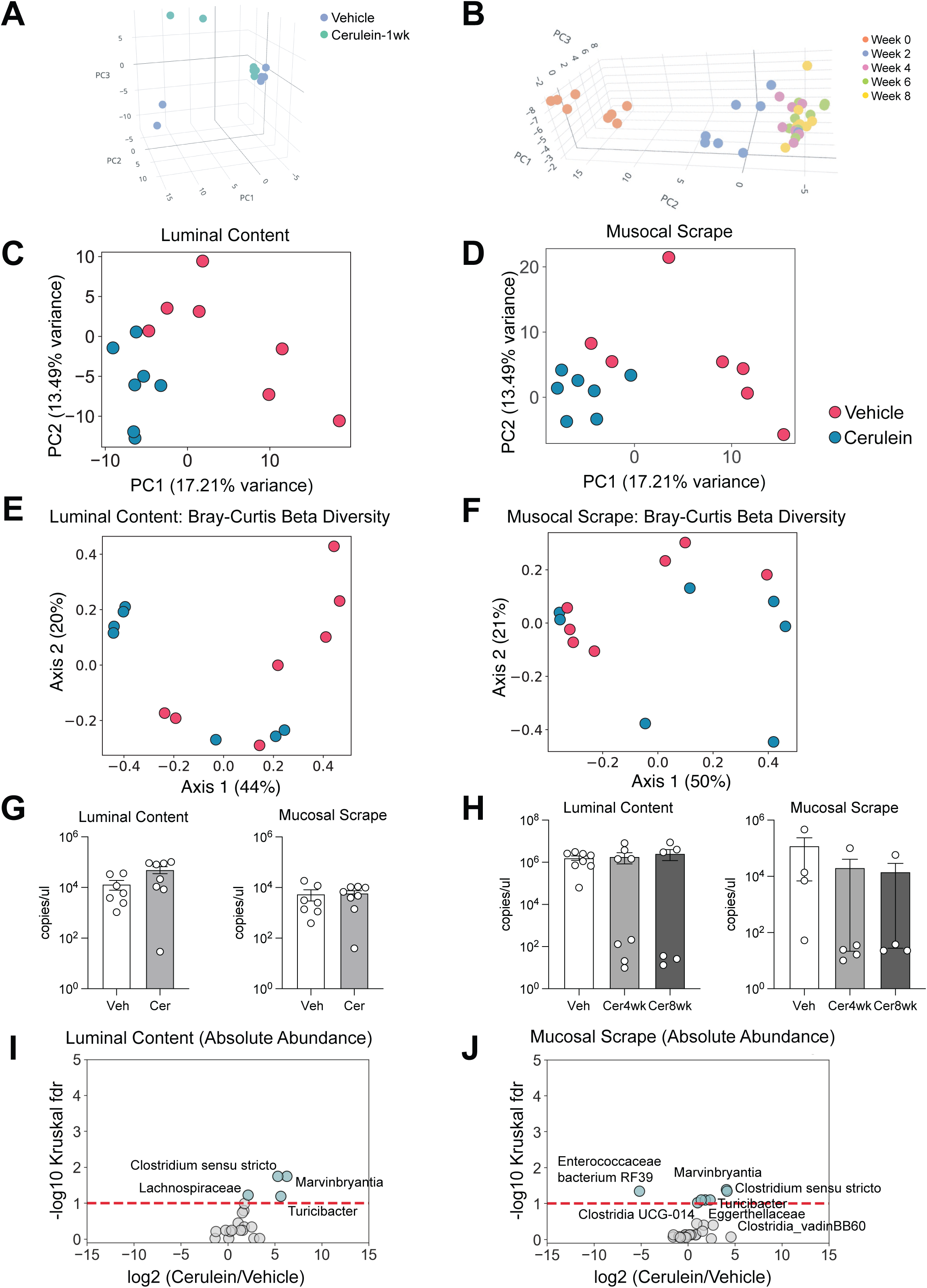
Cerulein induced pancreatitis modifies the gut microbiome. **A, B**. PCA analysis of fecal microbiome of mice treated with cerulein or vehicle for 1 week (**A**) or 4 or 8 weeks (**B**) (*n*=8 per group). **C, D**. PCA of CLR (centered log-ratio transformation) of duodenal luminal bacteria (**C**) and mucosa-associated bacteria (**D**) of mice treated with cerulein or vehicle for 1 week (absolute abundance) (*n*=8). **E, F.** Bray-Curtis beta-diversity analysis of duodenal luminal bacteria (absolute abundance) (**E**) and mucosa-associated bacteria (**F**) of mice treated with cerulein or vehicle for 1 week (*n*=8). **G, H.** Total microbial load in duodenal mucosa or lumen of mice treated with cerulein for 1 week (**G**) or 4 or 8 weeks (**H**) or vehicle control measured by quantitative 16S Q-PCR (*n*=8, samples with amount below detection threshold were excluded). **I, J.** Volcano plots of bacteria enriched or de-enriched in the duodenal lumen (**I**) or mucosa (**J**) of mice treated with cerulein or vehicle for 1 week (*n*=8).

